# A general theory for the dynamics of social populations: Within-group density dependence and between-group processes

**DOI:** 10.1101/2022.10.07.511281

**Authors:** Brian A. Lerch, Karen C. Abbott

**Author notes:** Open research statement*: Code to replicate analyses and figures has been uploaded to Zenodo. It can be accessed via https://zenodo.org/record/7063010#.Yx8Q0aHMJ3i.

## Abstract

Despite the importance of population structures throughout ecology, relatively little theoretical attention has been paid to understanding the implications of social groups for population dynamics. The dynamics of socially structured populations differ substantially from those of unstructured or metapopulation-structured populations, because social groups themselves may split, fuse, and compete. These “between-group processes” have been suggested to be important drivers of the dynamics of socially-structured populations, but no general theoretical framework exists that can handle various density-dependent between-group processes within a single model. Here, we develop a general framework for the dynamics of socially-structured populations that considers births, deaths, migration, group extinction, fissions, fusions, and between-group competition within a single model. Both logistic growth and an Allee effect are considered for within-group density dependence. We show that the effect of various between-group processes is mediated by their influence on the stable distribution of group sizes, with the ultimate impact on the population determined by the interaction between the altered group size distribution and within-group density dependence. The group level is important to the dynamics of the entire population, since it drives extinction risk, impacts population growth rates, and leads to the emergence of population-level density dependence (even if birth and death rates depend only on group size and not population size). We conclude with a series of case studies that illustrate different ways that age, sex, and class structure impact the dynamics of social populations. In sum, our results make clear the importance of within-group density dependence, between-group dynamics, and the interactions between them for the population dynamics of social species and provide a general, flexible framework for modeling social populations.

## 1 Introduction

Properly accounting for population structures is vital for understanding the dynamics of structured populations. The importance of population structures in shaping dynamics has been demonstrated in a wide range of contexts, such as metapopulation and metacommunitity ecology (Hanski, 1999; Leibold and Chase, 2017), epidemiology (Danon et al., 2011), and social evolution (Akçay, 2020). In any structured population, dynamics emerge from interactions that occur at a level that is below that of the entire population (Killingback and Doebeli, 1996; Molofsky et al., 1999; Bateman et al., 2018). Thus, to study structured populations, it must be determined whether and how processes occurring locally scale to the population level or else lead to new, emergent dynamics at the population level (Nowak and May, 1992; Lerch et al., 2018; Thompson et al., 2020). While strong theoretical foundations exist for understanding the dynamical feedbacks between populations and population structures in these diverse settings, the importance of social structures in shaping population dynamics remains far less studied. This is important because processes occurring at the level of social groups themselves (such as fissions, fusions, and competition between groups) are known to be key drivers of the dynamics of socially-structured populations and often have no analog in other contexts. As an example, compare the extinction risk of small social groups to small patches in metapopulations. In both cases, dispersal can be expected to lead to a “rescue effect”, where large patches/groups provide an influx of individuals to small or extinct patches/groups, buffering against extinction at the population level (Brown and Kodric-Brown, 1977; Eriksson et al., 2014). However, if small social groups suffer from increased competition with large groups, then the rescue effect may be overwhelmed by the actions of groups themselves. Small groups could, however, mitigate this by fusing with other groups, reducing their extinction risk. There is no analog for these latter group-level effects in the existing theory of populations structured by space or other factors. Therefore, while existing theory provides good intuition for some roles population structure plays in population dynamics, many aspects of the dynamics of social populations are rich for theoretical development.

Understanding socially-structured populations has important, real-world implications. For example, many social animals are also highly endangered, and properly accounting for social structures is required to make informed management decisions. Much of what we know about population dynamics in social animals comes from empirical work in these systems. Across taxa, between-group interactions can play a central role in the population dynamics of social species. For example, an increase in the number of groups in a mountain gorilla (*Gorilla beringei*) population led to a dramatic increase in violent intergroup encounters that drove a large decline in population growth, suggesting that the density of social groups (not individuals) regulates population density (Caillaud et al., 2020). This has also been suggested in lions (*Panthera leo*; Packer et al., 2005). Between-group competition has been shown to have important impacts on demographic rates in baboons (*Papio cynocephalus*; Markham et al., 2012), chimpanzees (*Pan troglodytes*; Lemoine et al., 2020), colobus monkeys (*Colobus guereza*; Harris, 2006), banded mongooses (*Mungos mungo*; Johnstone et al., 2020), and black-throated blue warblers (*Dendroica caerulescens*; Sillett et al., 2004). The fact that large groups are more likely to win such between-group conflicts (Majolo et al., 2020) can offset the costs of within-group competition, leading to intermediate group sizes being optimal (Cheney and Seyfarth, 1987; Markham et al., 2015).

More generally, intermediate group sizes may be optimal as a result of small groups having decreased growth rates (a group-level demographic Allee effect; Lerch et al., 2018). This has been observed in African wild dogs (*Lycaon pictus*; Rasmussen et al., 2008), vulturine guineafowl (*Acryllium vulturinum*; Papageorgiou and Farine, 2020), and social spiders (*Anelosimus eximius*; Avilés and Tufiño, 1998). Much attention has been paid to whether these density-dependent effects that decrease the growth rate of small groups will lead to small populations also having a decreased growth rate (a population-level demographic Allee effect). A consensus is building that this does not occur: within-group density dependence does not to scale to density dependence at the population level in multiple studies (Bateman et al., 2011; Woodroffe, 2011; Angulo et al., 2018), though population-level demographic Allee effects are possible in social species (Keynan and Ridley, 2016). Once again, between-group processes are key mediators of density-dependent effects (or the lack thereof) at the population level (Bateman et al., 2012). Most broadly, this occurs because group size is independent of population size (Angulo et al., 2013) due, for instance, to large groups acting as both source populations for small groups (Bateman et al., 2013) and drivers of new group formation (Woodroffe et al., 2020a). Empirical advancements for understanding the movement of individuals between groups are thus vital for a complete picture of social populations (McNutt, 1996; Maag et al., 2018; Behr et al., 2020; Woodroffe et al., 2020b).

Theory has been motivated by the aforementioned empirical trends. Most often, models have been developed to tackle specific questions. For example, theory has sought explanations for why within-group density dependence does not scale to the population level. Either social structure (Lerch et al., 2018) or group size heterogeneity (Angulo et al., 2018) can buffer against population-level demographic Allee effects arising from group-level demographic Allee effects. Known population-level Allee effects in social species are instead generated through other mechanisms that operate outside the level of the group. For example, if new group formation is stunted in small populations, a population-level demographic Allee effect can arise (Courchamp et al., 2000). Because sociality is often closely connected to behaviors that inherently involve sex or age structure, models of social species’ population dynamics often include such individual heterogeneity in addition to social structure. This improves realism, but means that isolating the effects of social structure requires additional work.

Given the focus on specific questions and systems, previously developed models tend to be limited in their scope through their use of functional forms tailored for a specific system and by considering only a narrow subset of between-group processes. Thus, there is a need for a general mathematical framework for studying the dynamics of social populations. The most general model to date is Bateman et al. (2018), which considers a socially-structured analog of matrix projection models (Cawell, 2006), where the number of groups of various sizes are tracked and transitions occur through births, deaths, and new group formation. The model of Bateman et al. (2018) is flexible, simple, and can readily be connected to empirical data. However, it is inherently linear (density-independent) at the population scale, meaning its asymptotic behavior is almost always exponential growth or decline. This makes it most appropriate for studying populations in an exponential phase of recovery or decline and the dynamics of group size distributions, rather than populations approaching a stable equilibrium, where the information provided by an exponential model is more limited. To study the dynamics of both the population and the social groups it comprises, a general framework that can account for population-level density dependence and non-linear processes occurring at the group level is needed.

To that end, we aim to develop a general theory for the dynamics of socially-structured populations that can account for a wide range of between-group processes. We adapt a theoretical framework for group selection (Simon, 2010; Simon et al., 2013) that has proven flexible for studying social evolution, for example in the contexts of cultural evolution (Henriques et al., 2019) and host-microbiome coevolution (van Vliet and Doebeli, 2019). These models are also a generalization of a continuous-time analog of Bateman et al. (2018) that allows for non-linearity and population regulation. Since they were developed for evolutionary questions, these models have never been analyzed thoroughly for their population dynamics. Here, we present these models in the context of population and behavioral ecology and provide a thorough analysis for the interplay of group and population dynamics in socially-structured populations. We begin by presenting a model that focuses on interactions occurring within and between social groups assuming all individuals are ecologically equivalent (i.e., lacking age-, sex-, or other population structure beyond social groups; see section 2.1 for a broad overview) and explaining what information can be obtained from various implementations of the model (section 3). Results from this model clarify the role of various between-group processes play in the dynamics of social populations (section 4). We then generalize the model to permit individuals to play ecologically distinct roles and provide a series of case studies demonstrating the importance of individual heterogeneity (age, sex, and class structure) in the population dynamics of social species (section 5). Broadly, we find the stable distribution of group sizes is central to understanding population dynamics. The effect of various between-group processes can be inferred from the way that they alter the group size distribution (with the ultimate impact being determined by an interaction with within-group density dependence). We find that socially-structured populations are at a heightened extinction risk relative to unstructured populations of the same size. Further, without processes that are density-dependent at the population level, populations either grow or decline exponentially (regardless of density dependence within social groups). Processes occurring at the group level whose rates depend on the density of social groups can themselves regulate populations, even when individual birth and death rates do not depend explicitly on the population density.

## 2 Model: Ecologically identical individuals

We now present the biological assumptions of our initial model, illustrating its simplest form. A summary of parameters, variables, functions, and other notation is provided in Table 1.

**Table 1:**
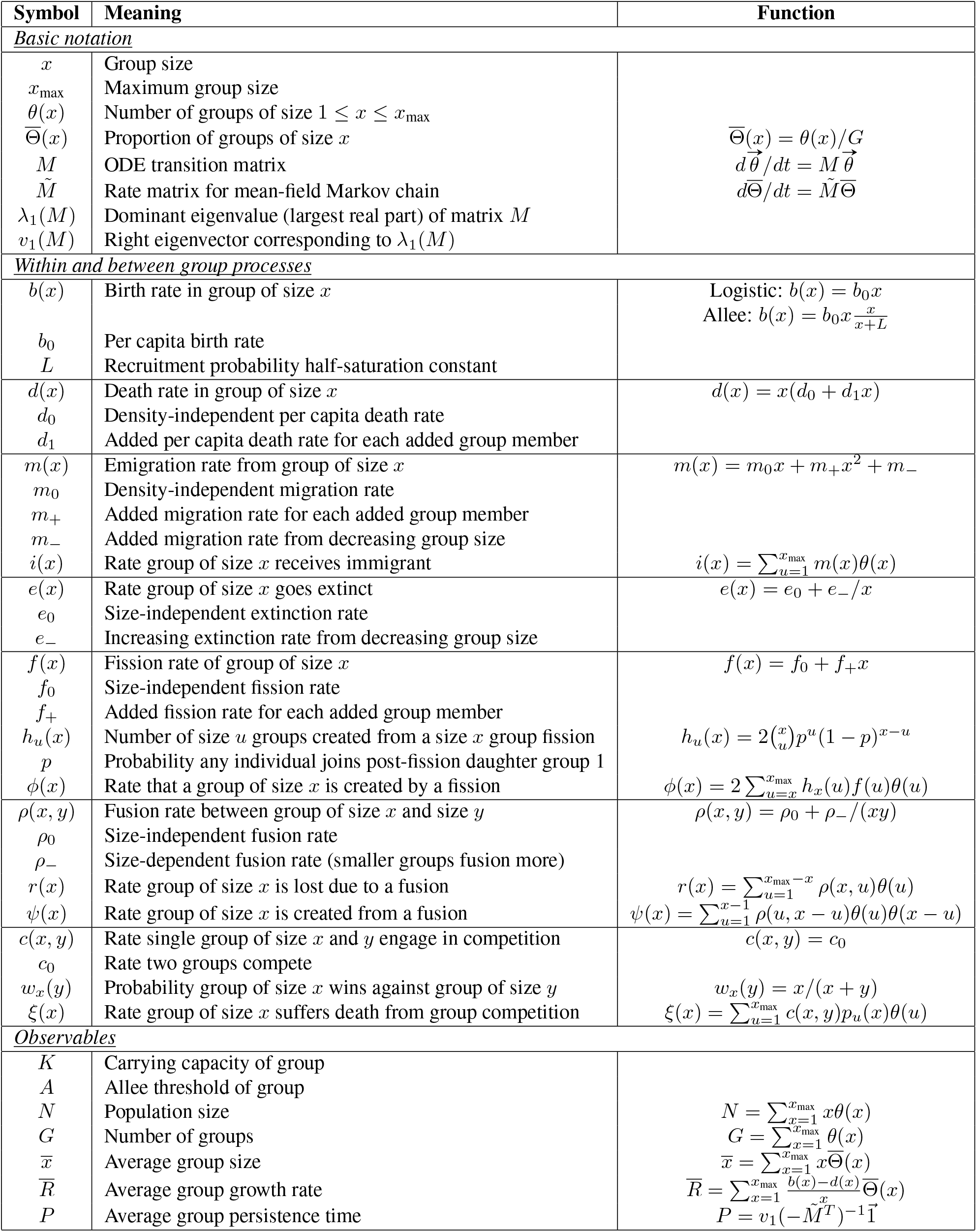
Summary of model parameters, functions, and variables.

### 2.1 Overview

We assume that all individuals in the population are equivalent (an assumption that we will relax below in section 5). The population consists of social groups of various sizes 1 ≤ *x* ≤ *x*_max_. At the population level, our model tracks the number of groups of size *x*, *θ*(*x*). For completeness, we provide a description of the model here, noting the close accord with previous literature (Simon, 2010; Simon et al., 2013). Ultimately, the dynamics of our model are governed by the way in which various processes (e.g., births, deaths, fissions, migration) alter the group size distribution. For example, a birth in a group of size *x* causes the group to transition to size *x* + 1. We begin by describing the underlying stochastic processes of our model (e.g., the biological features of populations that we consider): births, deaths, and five kinds of “between-group processes,” since these processes are equivalent in all versions of the model. Only after describing these underlying processes in the sections 2.2 and 2.3, do we formalize the way in which the model tracks population dynamics in section 3.

Our primary goals are i) methodological (to provide a general model of the dynamics of social populations) and ii) biological (to describe the qualitative influence of between-group processes on the dynamics of social populations). Due to the former goal, portions of this paper are somewhat technical for readers that simply want the biological conclusions. Such readers can reference Fig. 1 for a broad overview of our model and skip to the results (section 4) without missing biological conclusions. Readers that wish to see how we mathematically describe various social processes without being bogged down in technical details should read the remainder of section 2 then skip to the results (section 4). Section 3 provides the most technical details of our model and analysis.

**Figure 1:**
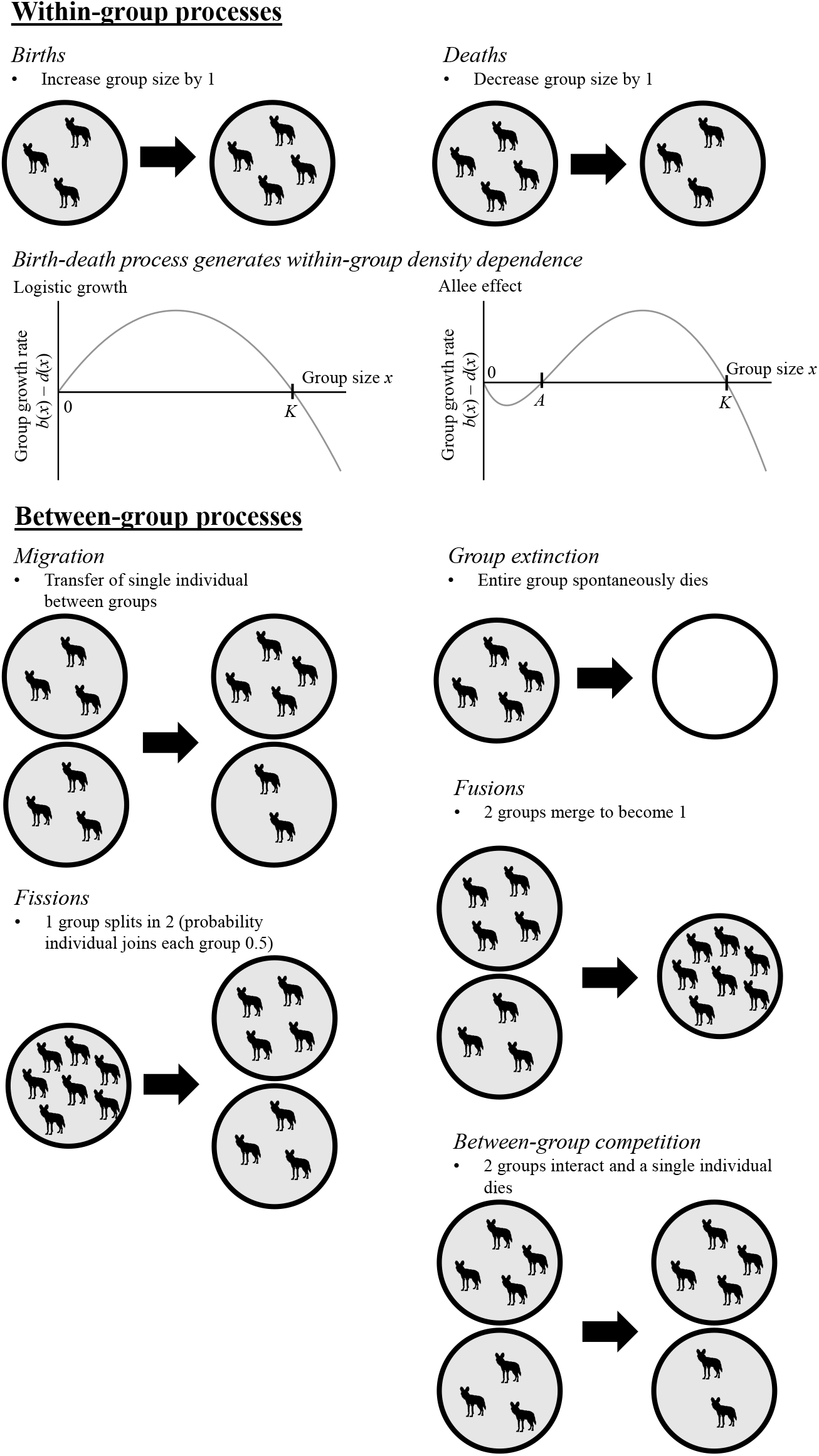
Conceptual overview of our model. We always consider exactly two within-group processes (births and deaths) that result in either negative or positive density dependence (logistic growth or an Allee effect) within the group. We also consider five between-group processes that involve the transfer of individuals between groups or group-level influences on group dynamics.

### 2.2 Within-group processes

We refer to births and deaths as “within-group processes” as these involve solely individuals within a single group. Births and deaths are the most basic processes that we model and to ensure biological realism we always permit each to occur with non-negative rates. We consider two types of density-dependence: logistic growth and an Allee effect (Fig. 1).

#### 2.2.1 Logistic growth

Logistic growth within groups signifies inherently competitive interactions within groups, as it results in the per capita growth rate declining linearly with group size. To produce logistic growth, we assume a constant per capita birth rate *b*_0_ such that the overall birth rate in a group of size *x*, *b*(*x*), is

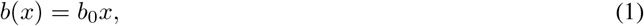

with the added assumption that *b*(*x*_max_) = 0 to bound group size at *x*_max_. For death rates *d*, we assume that the per capita death rate increases linearly with group size such that

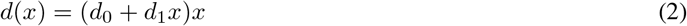

is the overall death rate in a group of size *x*. Here, *d*_0_ is the per capita death rate in a (hypothetical) group of size 0 and each additional individual in the group results in an increase of *d*_1_ in the per capita death rate. The stable group size *K* for logistic growth can be found by setting *b*(*x*) − *d*(*x*) = 0 and solving for *x*. With the birth-death process given by Eqn. 1 and 2, 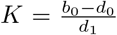. We refer to *K* as the carrying capacity of groups. In practice, we set *b*_0_, *d*_0_, and *K* and solve for *d*_1_ rather than fixing *d*_1_ in advance.

#### 2.2.2 Allee effect

We also consider a group-level demographic Allee effect (where the expected group growth rate increases with group size in small groups; Lerch et al., 2018), often cited as a signature of cooperative interactions. To model the Allee effect, we use the birth-death process from Dennis (2002). Now, the birth rate in a group of size *x* is given by

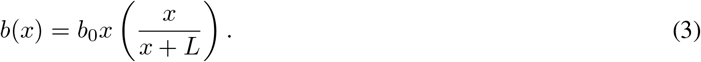

*b*_0_ is again the intrinsic per capita birth rate. Contrasting with Dennis (2002), we interpret the term in the parentheses to be the probability that any birth is recruited into the population. This recruitment probability saturates with increasing *x* with *L* as the half-saturation constant (i.e., the group size in which half of all births are expected to be recruited).

The death rate for the Allee effect remains unchanged from logistic growth (Eqn. 2). The birth-death process given by Eqn. 3 and 2 predicts two equilibrium group sizes: the smaller is unstable and we refer to it as the Allee threshold *A*, while the larger is stable and we refer to it as the carrying capacity *K*. For group sizes smaller than the Allee threshold or larger than the carrying capacity (*x* < *A* or *x* > *K*), the expected group growth rate is negative. Only for group sizes in between the Allee threshold and carrying capacity (*A* < *x* < *K*) is the expected group growth rate positive. To facilitate both biological interpretation and comparison with logistic growth, in practice we fix *b*_0_, *d*_0_, *A*, and *K* in advance and solve for *d*_1_ and *L* using the system of equations

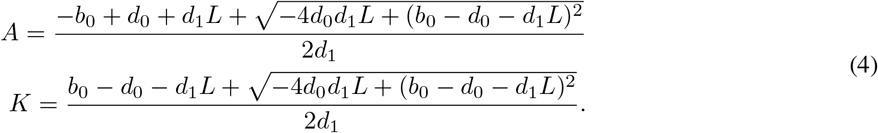

### 2.3 Between-group processes

Between-group processes are the core feature of our model and central to understanding population dynamics in social species. We define between-group processes to mean any event that involves either i) individuals from more than one group (e.g., fusions or group competition), ii) a change in the size of more than one group (e.g., migration), or iii) a spontaneous change in the number of groups (e.g., fissions or group extinctions). This characterization highlights the fact that groups themselves are dynamic entities that influence population dynamics. We note that our definition differs from “group-level events” as defined by Simon et al. (2013) which, for example, excludes migration. Rather than quarreling with the characterization in Simon et al. (2013), differing definitions reflect our different focus (here population dynamics, there group selection). We explicitly formulate and analyze five between-group processes: migration, group extinctions, fissions, fusions, and group competition (Fig. 1). Other between-group processes that may influence dynamics in specific systems can readily be incorporated into this modeling framework. Below, we describe each between-group process mathematically.

#### 2.3.1 Migration

Migration is the transfer of an individual between two groups (Fig. 1). We assume that the total rate that any individual emigrates from a group of size *x* is

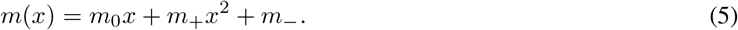

*m*_0_ accounts for a constant per capita emigration rate, *m*_+_ the added per capita emigration rate for each additional group members (individuals being more likely to emigrate from large groups), and *m*_−_ for individuals being more likely to emigrate from small groups. We assume that once an individual emigrates, it instantaneously joins a new group randomly, provided it can join without causing the group to exceed the maximum size. Then, the rate at which a given group of size *x* acquires an immigrant is

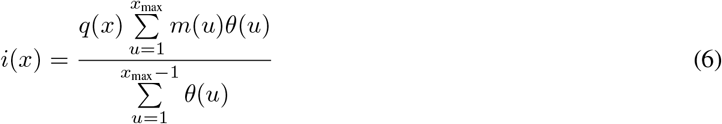

with

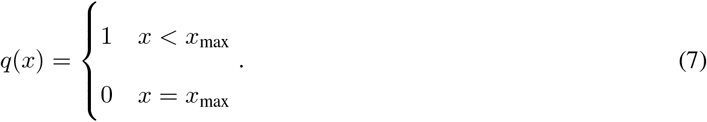

The immigration rate into a group of size *x* (Eq. 6) is given by the probability that the chosen group accepts the immigrant *q*(*x*) times the rate at which individuals emigrate (the sum in the numerator) divided by the total number of groups that could accept an immigrant (i.e., the probability the focal group is chosen; the denominator). Here, we assume that any group that is not already full will accept immigrants, though more complex functions could readily be incorporated into how *q*(*x*) is defined in Eq. 7. One could also use *q*(*x*) to control how likely a dispersing individual is to attempt to immigrate into groups of various sizes (e.g., a bias for immigrating into intermediate-size groups). However, since only the net flux of migration determines its qualitative influence, and because we already consider per capita emigration rates changing with group size, we do not consider the case of immigration biased by group size here.

#### 2.3.2 Extinction

Though groups in our model can go extinct due to demographic stochasticity, there may be additional mechanisms that cause groups to abruptly go extinct due to processes other than the deaths already accounted for elsewhere in the model (e.g., disease). These extinction events are modeled by instantaneously causing the death of all individuals within a group (Fig. 1). We assume that such stochastic group extinctions occur at rate

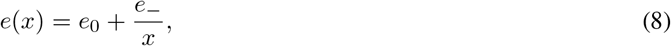

where *e*_0_ is the extinction rate regardless of group size and the *e*_−_ term accounts for smaller groups going extinct more frequently.

#### 2.3.3 Fission

We next model fissions, whereby a single group splits into two groups (Fig. 1). We assume the rate at which a single group of size *x* fissions is

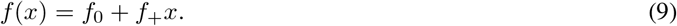

Here, *f*_0_ is the fission rate independent of group size and *f*_+_ is the additional rate of fissioning resulting from each additional individual (i.e., if *f*_+_ > 0, larger groups fission more frequently). We assume that the probability of a specific individual joining post-fission group 1 (an arbitrary but fixed group) is *p* and that individual decisions are independent. Then, the distribution of post-fission group sizes is binomially distributed with *x* trials and success probability *p*. For species that deviate from these assumptions, other distributions could be used. Under the binomial assumption, the expected number of groups of size *u* produced from a fission of a group of size *x* (hereafter, the “fissioning density”) is (Simon, 2010)

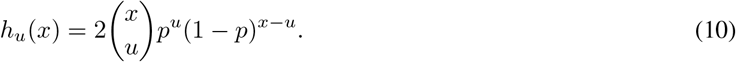

The factor of 2 reflects that, due to the total number of individuals being conserved through the fission, assigning *u* or *x − u* individuals to post-fission group 1 both result in one new group of size *u*. Eqn. 10 is the expected number of groups of size *u* created given that a group of size *x* fissions. Then, in total, the rate at which a group of size *u* is created from a fission is

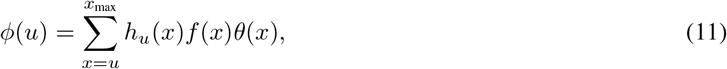

where *f* (*x*)*θ*(*x*) is the rate at which there is a group of size *x* that fissions and the sum is over all group sizes that could fission and produce a group of size *u*. Because the fission rate of a single group *f* does not depend on the number of groups, fissions are density-independent at the population level (that is, *f* is a function only of *x*, not *θ*(*x*)).

This means that the rate groups form due to fissions (*ϕ*) depends linearly on the number of groups of various sizes, *θ*(*x*), and thus the standard tools for analyzing linear models may be used even when groups fission at a non-zero rate. Throughout we restrict our analysis to the case where *p* = 1/2 (that is, individuals tend to form approximately equally-sized groups post fission), though clearly other values of *p* could be used for species with uneven post-fission group sizes.

#### 2.3.4 Fusion

The opposite of a group fission is the fusion of two groups. A fusion occurs when two groups come into contact and all individuals merge to become a single group (Fig. 1). The rate at which a single group of size *x* fuses with a single group of size *y* is given by

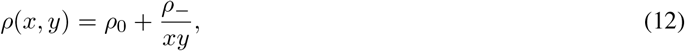

for *x*, *y* > 0 and *x* + *y* ≤ *x*_max_. Here, *ρ*_0_ is the fusion rate independent of group size and *ρ*_−_ accounts for increasing fusion rates of small groups. In contrast with processes we have considered thus far, we assume that fusions become more frequent with more groups in the population due to increased interaction rates. In particular, we assume that groups fuse in proportion to the rate at which they contact one another through mass action assumptions. Thus, at the population level, the total fusion rate between any group of size *x* and any group of size *y* is *ρ*(*x*, *y*)*θ*(*x*)*θ*(*y*). Then, the per-group rate at which a group of size *x* is lost due to a fusion *r*(*x*) is the sum over all possible fusions of groups of size *x* such that

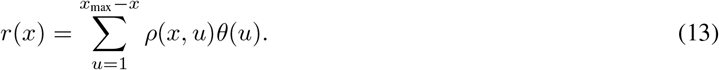

Note that although *ρ*(*x*, *y*) is technically defined for any combination of pre-fusion group sizes, the upper limit in this summation precludes fusions that would result in a new group exceeding *x*_max_ individuals. Likewise, the total rate at which a group of size *x* is created due to a fusion *ψ*(*x*) is the sum over the rates of all fusions that can generate a group of size *x*. Thus,

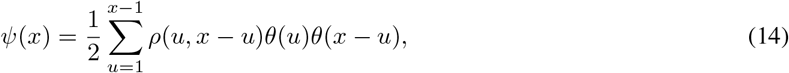

where the factor 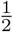 accounts for the fact that each fusion between two groups only generates a single group.

#### 2.3.5 Group competition

Much like fusions, we assume that group competition occurs more frequently when there are more groups in the population. We consider that a group of size *x* interacts with a group of size *y* at rate *c*(*x*, *y*). Thus, total rate of competition between any group of size *x* and any group of size *y* is *c*(*x*, *y*)*θ*(*x*)*θ*(*y*). We assume that this rate is constant (*c*(*x*, *y*) = *c*_0_), though more complicated functions could be considered (including asymmetrical functions that model the case where only one of the two groups is affected by the competition). Upon competing, we assume that exactly one group will “win” the competition, larger groups are more likely to win, and losing the competition results in the death of a single group member. In other words, group competition behaves exactly as an added death rate to the losing group (though other impacts of competition can be readily incorporated and we will briefly consider an alternative formulation where competition results in the extinction of an entire group in the results). The probability that a group of size *x* wins competition with a group of size *y* is assumed to be

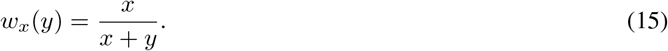

Under these assumptions the total rate of death in a given group of size *x* due to group competition is

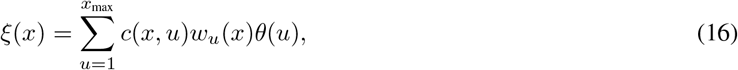

where the sum accounts for rate of competition (and the probability of winning) against all other groups in the population.

## 3 Model forms and analyses

Above, we presented the biology underlying our model and how we represent the biology with equations. Now, we turn to mapping biological features into various modeling frameworks. We include a multitude of frameworks because each provide us with slightly different information about population dynamics and thus is best suited to answering different sets of questions about a system’s ecology. We will structure this section by first explaining a modeling framework that we consider (i.e., a way to generate dynamics from the biological processes above) and then explaining what analyses can be carried out on the given framework and what information it provides. It is generally impractical to track the dynamics of every single group in a population, so tractable approaches involve a tradeoff between understanding the entire population at the expense of knowing the fates of individual groups, and vice versa. The best modeling strategy depends on what questions we wish to answer, as illustrated below.

### 3.1 Population-centered Markov chain

We begin with the most fundamental model to track the dynamics of a social population under the biological assumptions detailed above: the population-centered Markov chain. We use “population-centered” to stress that the sizes of an entire collection of social groups (not just one) are being tracked. This model is best suited for questions that require accounting for demographic stochasticity.

#### 3.1.1 Formulation

Model dynamics occur as a result of various stochastic processes that occur within and between groups (e.g., births, deaths, fissions, migration, etc.). We assume that each obeys a Poisson process; that is, the waiting time between events is exponentially distributed and thus the dynamics follow a continuous-time Markov chain. The various processes cause a change in the distribution of group sizes. For example, a birth in a group of size *x −* 1 results in one fewer group of size *x* − 1 (*θ*(*x* − 1) → *θ*(*x* − 1) *−* 1) and one more group of size *x* (*θ*(*x*) → *θ*(*x*) + 1). Thus, the state of the system is given by {*θ*(*x*): 1 ≤ *x* ≤ *x*_max_}. Rates and the resulting state transitions for the population-centered Markov chain are summarized in Table 2.

**Table 2:** State transitions in the population-centered Markov chain.

| Process | Transition(s) | Rate |
| --- | --- | --- |
| Birth | $\theta(x) \rightarrow \theta(x) - 1$<br>$\theta(x + 1) \rightarrow \theta(x + 1) + 1$ | $b(x)\theta(x)$ |
| Death | $\theta(x) \rightarrow \theta(x) - 1$<br>$\theta(x - 1) \rightarrow \theta(x - 1) + 1$ | $d(x)\theta(x)$ |
| Migration | $\theta(x) \rightarrow \theta(x) - 1$<br>$\theta(y) \rightarrow \theta(y) + 1$ | $m(x)\theta(x)\theta(y)/(G - \theta(x_{\max}))$ |
| Extinction | $\theta(x) \rightarrow \theta(x) - 1$ | $e(x)\theta(x)$ |
| Fission | $\theta(x) \rightarrow \theta(x) - 1$<br>$\theta(u) \rightarrow \theta(u) + 1$<br>$\theta(x - u) \rightarrow \theta(x - u) + 1$ | $h_u(x)f(x)\theta(x)$ |
| Fusion | $\theta(x) \rightarrow \theta(x) - 1$<br>$\theta(y) \rightarrow \theta(y) + 1$<br>$\theta(x + y) \rightarrow \theta(x + y) + 1$ | $\rho(x, y)\theta(x)\theta(y)$ |
| Group competition | $\theta(x) \rightarrow \theta(x) - 1$<br>$\theta(x - 1) \rightarrow \theta(x - 1) + 1$ | $c(x, y)w_y(x)\theta(x)\theta(y)$ |

Without a maximum number of groups of a given size (a quantity referred to as *κ*), the number of states is infinite. Even with a finite *κ*, the number of states is extremely large 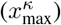. This prevents us from writing down the Markov chain’s rate matrix for analytical treatment. We must thus resort to simulation; we use Gillespie’s stochastic simulation algorithm (Gillespie, 1977).

#### 3.1.2 Analysis: Extinction probability

The inclusion of demographic stochasticity in the population-centered Markov chain model is particularly useful considering that the relatively small size of many social groups makes them especially susceptible to the effects of demographic stochasticity. The population-centered Markov chain is the only stochastic model form that we consider, and thus only it can be used to assess extinction risk. Unless the population displays unbounded growth, it will go extinct with probability 1 given infinite time. We simulate 1000 trajectories from the population-centered Markov chain for 2000 time units and compute the fraction of trajectories that have gone extinct by time *t* (for all 0 < *t* < 2000) to approximate the probability of extinction by time *t*.

### 3.2 Population-centered deterministic model

We now consider the deterministic limit of the population-centered Markov chain as the number of groups becomes infinite. The population-centered deterministic model is in many ways simpler to work with than the corresponding Markov chain as it is computationally efficient and can be easily interpreted. This model is well-suited for gaining a general understanding of patterns of population growth.

#### 3.2.1 Formulation

By taking the limit as the number of groups goes to infinity appropriately, our population-centered Markov chain converges to a system of *x*_max_ ordinary differential equations (ODEs) (Kurtz, 1970; Puhalskii and Simon, 2012). The population-centered deterministic model can be concisely written as

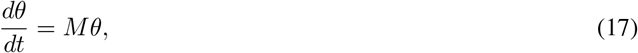

where *θ* = (*θ*(1), *θ*(2), …, *θ*(*x*_max_))^*T*^ is the vector for the number (technically, density) of groups of size 1, 2, …, *x*_max_, respectively. The *x*_max_ *× x*_max_ matrix *M* is the rate matrix for the system of ODEs that describes changes in group size. The element in the *i*th row and *j*th column of *M*, denoted *M* (*i*, *j*) is (for *i* ≠ *j*) the rate at which a group of size *i* is created per group of size *j*. Diagonal elements *M* (*i*, *i*) are given by negative the rate at which groups of size *i* are lost. With non-linear between-group processes, *M* will itself be a function of *θ*. It is thus easier to write out the general form of the differential equation for the number of groups of an arbitrary size *x*,

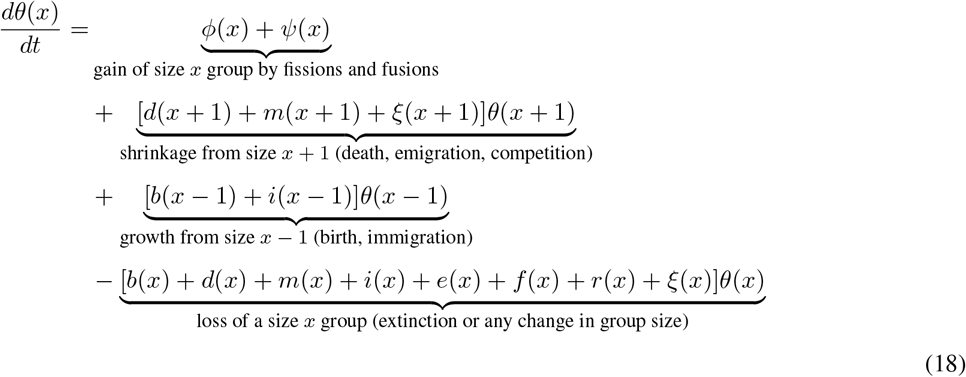

for 1 ≤ *x ≤ x*_max_ with the boundary conditions *θ*(0) = *θ*(*x*_max_ + 1). Here, the first row of the right-hand side accounts for the generation of groups of size *x* through processes that can cause large changes in group size – that is, group formation through fissions and fusions. The second row accounts for processes that cause groups of size *x* + 1 to lose a single individual (deaths, emigration, and group competition). The third row accounts for processes that cause groups of size *x* − 1 to gain a single individual (births and immigration). The fourth row accounts for all processes that cause the loss of a group of size *x*, whether that leads to the transition to a group of another size or the spontaneous extinction of the group. Appropriately collecting coefficients to *θ*(*j*) for the equation with *x* = *i* allows one to write *M* (*i*, *j*). Note that whenever non-linear between-group processes are included, *M* is not unique (because terms of the form *θ*(*i*)*θ*(*j*) could appear either in the *j*th column or the *i*th column); however, the resulting dynamic equations do not depend on the specific choice of *M* from among these possibilities. Note that it has also been conjectured that taking different limits result in a deterministic model of a single, partial differential equation (Simon et al., 2013) but we do not consider such a model here.

#### 3.2.2 Analysis: Group size distribution

Whenever our population-centered deterministic model is linear, the number of groups will shrink or grow exponentially at rate given by the eigenvalue with largest real part, which we call *λ*_1_, of the transition matrix, *M*. In spite of exponential growth, the relative frequency of each group size approaches a constant value. In the context of our model, this is a (unique) stable distribution of group sizes that will remain fixed as the population grows or shrinks (we refer to this as the “group size distribution” and denote it by 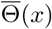). Whenever the population-centered deterministic model is linear, the group size distribution can be computed as the right eigenvector associated with the largest eigenvalue (*v*_1_). If the population-centered deterministic model is non-linear, then the group size distribution is not guaranteed to equilibrate (though we find that in our model it does for the functions considered) nor be unique. In this case, we compute the group size distribution for given initial conditions by numerically integrating Eqn.18.

We chose to present the group size distribution analysis here since it is most straightforward using the population-centered deterministic model, but it is worth acknowledging that the population-centered Markov chain can be also be studied for its group size distribution. However, as we noted above, any population that is subject to stochasticity and is not growing exponentially will eventually go extinct; as such, the stationary distribution of the Markov chain is extinction of the entire population. However, prior to extinction, there is a quasi-stationary distribution of group sizes of the Markov chain (conditioned upon the population not being extinct). The group size distribution is the expected value of this quasi-stationary distribution.

#### 3.2.3 Analysis: Density dependence and growth rates

As explained above, when the model is linear in the number of groups, the expected rate of growth of the number of groups is given by the largest eigenvalue of *M*, *λ*_1_. Since losing a group, on average, results in a loss of an average group size 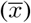 number of individuals, this means that we can compute the per capita population growth rate as 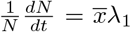. This equation makes plain how density dependence cannot arise at the population level, regardless of the group-level density dependence, as long as the population-centered deterministic model is linear. The above equation no longer holds when there are non-linearities, but there are still straightforward ways of assessing the per capita population growth rate. Notice that

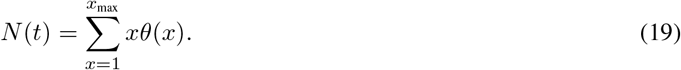

Differentiating, this implies that

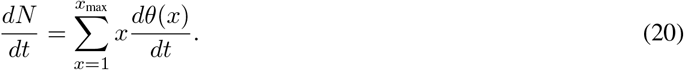

The term 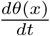 has already been specified as the population-centered deterministic model (Eqn. 18). Plugging in and simplifying yields

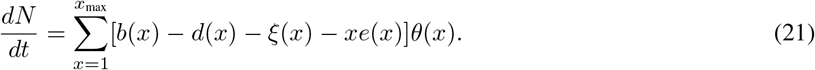

Note that this equation contains only those processes that add (birth) or subtract (death, competition, or extinction) individuals, as processes that simply move individuals among groups will not change the total population size and therefore cancel when we take the sum over all group sizes. Assuming the existence of a stationary group size distribution and using the identity that in a population with *G* groups, 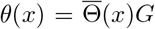 and 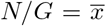, the per capita population growth rate can be written as

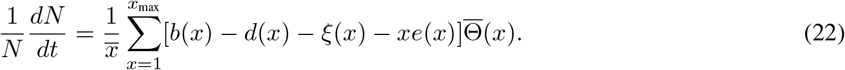

Thus, the population growth rate is the rate at which individuals are born in the entire population minus the rate individuals die or are lost from groups of any size going extinct.

One important implication of Eqn. 22 is that even though many between-group process (all except extinction and group competition) conserve the total number of individuals in the population, they can still affect the population growth rate via their effect on the group size distribution. Indeed, changing birth, death, and group competition rates can also influence the group size distribution and thus the population growth rate indirectly, in addition to their direct effects. The second important implication of Eqn. 22 is that social populations will grow in a density-independent manner unless the average group size or group size distribution changes with population size. That is, only functions of group size (*x*), not population size (*N*), appear within the square brackets in Eqn. 22, reflecting how the direct density dependence experienced by social species occurs at the level of the group. This means that the only way for a between-group process to generate population-level density dependence is if it causes the group size distribution to be a function of the overall population size. Below, we ask when this occurs.

In addition to population growth, it is often desirable to understand the pattern of group growth. We use the average per capita group growth rate across all groups in the population,

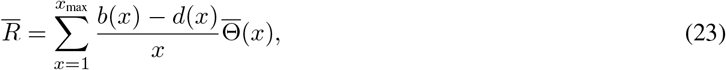

as a measure of within-group population growth. Since between-group processes change the group size distribution, this allows us to assess how the average rate of growth that individuals experience within their social groups changes as a result of between-group processes.

#### 3.2.4 Analysis: Reproductive value

A well-known result from matrix projection models is that the left eigenvector associated with the dominant eigenvalue of the population projection matrix gives the reproductive value of the various states (i.e., the contribution of each state to population growth; Cawell, 2006). We can perform the same analysis on our model by finding the left eigenvector (eigenvector of the transpose of *M*) associated with the dominant eigenvalue *λ*_1_. Care must be taken in interpreting reproductive value in this setting: it is the contribution of groups of various sizes to population growth (Bateman et al., 2018). Following Bateman et al. (2018), we rescale the reproductive value to be relative to individuals breeding independently (a group of size 1), by dividing each term by the first entry of the left eigenvector. Thus, as an example, a reproductive value of 2 for a group of size *x* means that groups of size *x* contribute twice the amount to population growth as individuals breeding independently.

### 3.3 Group-centered Markov chain

Above, we have focused on models that describe changes in the entire population. We have seen what information can be gained by considering all groups at once, but in doing so, we lose information about the trajectory of any one group through time. To better understand the dynamics of one group, we must take a “group-centered” approach that explicitly follows one group through time. We refer to this as the group-centered Markov chain. This model is best suited for questions involving the fate or trajectory of a single group.

#### 3.3.1 Formulation

The group-centered Markov chain is closely related to the population-centered models and the same tools as presented above can be used to study this scenario (albeit with slightly different interpretations). Let 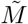 be the rate matrix describing transitions in the size of a single group. We highlight special cases for this model below.

##### Baseline: Without between-group processes

In the case with only births and deaths (no between-group processes), the dynamics of a single group obey a birth-death Markov chain. Here, deaths are denoted on the superdiagonal and births on the subdiagonal of the rate matrix. Thus, the reduced rate matrix 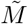 for the Markov chain describing transitions of this single group (i.e., the rate matrix removing the absorbing state of group extinction) is equivalent to the rate matrix of the population-centered Markov chain 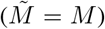. This implies that the population-centered model of numerous groups without between-group processes is ergodic (the distribution across groups at one time is the same as the distribution through time of one group).

##### Migration

Because immigration is non-linear and depends on the states of other groups in the population, additional assumptions must be made to model a single group experiencing immigration. We do so by assuming that there are 25 groups total in the population with distribution numerically calculated as 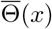 to set the immigration rate. The effect of emigration is equivalent to a death and the effect of immigration equivalent to a birth.

##### Fission

The dynamics of a single group that fissions at rate *f* (*x*) is a straightforward modification of the population-centered model. We make the simple assumption that when a group fissions a randomly chosen post-fission group retains the identity of the original group. Thus, rather than using the fissioning density to determine the post-fission group size, we simply use the binomial probability mass function (PMF).

##### Fusion

As with migration, fusions are non-linear and depend on the states of other groups in the population. We assume that the external population is made up of 25 groups that follow the group size distribution without between-group processes, which also gives the probability distribution for the focal groups initial size. Fusions involving the focal group are then modeled by adding the individuals from a randomly selected group from this distribution to the focal group (i.e., the focal group still “exists” after the fusion). The focal group’s size thus becomes larger in discrete jumps as a result of fusions.

#### 3.3.2 Analysis: Distribution of single group’s size through time

Because groups are not indexed in any way in the population-centered models, we can only observe what occurs in a single group through time by studying the group-centered Markov chain. The eigenvector associated with the dominant eigenvalue of the rate matrix 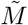 gives the distribution of a single group’s size through time. In contrast to the group size distribution described above (the relative frequencies of group sizes found in the population at a snapshot in time), this distribution follows a single group through time and computes the probability (conditioned on not going extinct) that a single group can be found at various sizes. This snapshot of a single group through time may or may not be equivalent to the snapshot of all groups at a single point in time. Whether these two snapshots are equivalent provides information about the relationship between the population-centered and group-centered models. The distribution of a single group’s size through time, in particular, is informative for understanding the influence of between-group processes on individual groups.

#### 3.3.3 Analysis: Group persistence time

The second piece of information that can be extracted from the group-centered Markov chain is the expected persistence time (mean time to extinction) of a single group. Assuming that the probability the group initially has *x* individuals is given by the group size distribution, the mean persistence time *P* of the group is (Durrett, 2016)

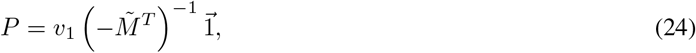

where *v*_1_ is the group size distribution as a column vector, the operator ^*T*^ indicates the transpose, the operator ^−1^ the inverse, and 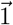 is the column vector of *x*_max_ 1s.

### 3.4 Two-variable model

Even our simplest model, the population-centered deterministic model, is largely analytically intractable. As a final model, we consider a simplification where we track only the number of groups *G* and the total population size *N*. We refer to this toy model as the “two-variable model” since it consists of only two state variables (compared to *x*_max_ variables for the population-centered deterministic model).

#### 3.4.1 Formulation

By condensing our model into two state variables, we lose all information about the distribution of group sizes. As such, we assume that all vital rates are functions of the average group size 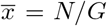. By losing information about the group size distribution, groups also do not go extinct naturally by experiencing a death when they consist of only one individual. As such, we assume that groups are lost at rate *ℓ*. Note that this is meant to capture the loss of groups due to demographic stochasticity (rather than spontaneous, stochastic extinction events). Thus, to make the two-variable model as comparable as possible to the population-centered deterministic model, we will use 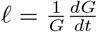 with 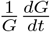 calculated as the long-run rate (*λ*_1_) from the population-centered deterministic model without between-group processes. We consider a few special cases for the two variable model below.

##### Baseline: Without between-group processes

We begin by showing the specific form for equations without between-group processes. Dynamics are given by

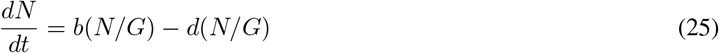

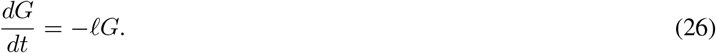

This is equivalent to assuming that the population consists of *G* groups that are all of the same size where groups go extinct at a constant per-group rate. Recall that we parameterize our model such that *ℓ* should be interpreted as the analog to extinction due to demographic stochasticity, not as the between-group process described above.

##### Migration

Migration between groups conserves both the population size and the number of groups (since there are no groups with only 1 individual that go extinct due to migration). As such, the two-variable model with migration is identical to the baseline two-variable model. This demonstrates that any effect due to migration from the other models relies upon group size heterogeneity.

##### Fission

Since fissions conserve the total population size, Eq. 25 does not change for the case of fissions. Fissions simply produce new groups at rate *f*_0_. As such the dynamics for number of groups (cf Eq. 26) becomes

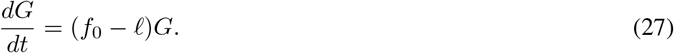

##### Fusion

Much like the case of fissions, fusions conserve the total population size and thus Eq. 25 does not change with fusions. Instead, the dynamics for number of groups (cf Eq. 26) now obeys

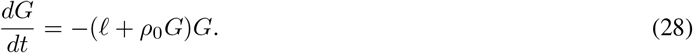

##### Group competition

In contrast to fissions and fusions, group competition conserves the number of groups, but not the number of individuals. Thus, dynamics for the number of groups obeys Eq. 26 and the dynamics for the population size follows

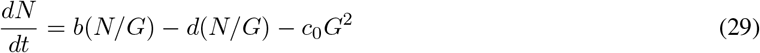

#### 3.4.2 Analysis: Role of group size heterogeneity

The two-variable model can be analyzed for its growth rates much like the population-centered deterministic model, but the model’s simplicity makes the results of such analyses rather limited. Its primary value instead is to demonstrate the role that group size heterogeneity plays in shaping the population dynamics of social species (argued to be a key driver of dynamics elsewhere; Angulo et al., 2013, 2018). If the two-variable model produces results that are qualitatively similar to the population-centered deterministic model, then group size heterogeneity is not the ultimate driver of model dynamics. Notably, this does not rule out group size heterogeneity playing some role, since *ℓ* is set to match a model with group size heterogeneity. However, it does demonstrate that it is not the heterogeneity *per se* that is driving dynamics (as similar results are obtained without group size heterogeneity).

## 4 Results: Ecologically identical individuals

We now present results from the models described above. Recall that the purpose of including different model forms is that each provides us with a different set of information about a system’s dynamics. To establish baseline expectations, we begin by considering the case with no between-group processes (section 4.1). This is equivalent to a case of metapopulations with no migration and, though perhaps not directly relevant to any real system, still informative for building intuition about the dynamics of structured populations. Next, we consider the role of each between-group process in isolation (section 4.2). In the previous section, we saw that even if they do not directly change the number of individuals in the group, between-group processes still can influence population dynamics by altering the group size distribution. We begin with migration and extinction as these cases are analogous to many results from metapopulation theory (though our interpretation for social populations is sometimes new). Next, we move on to between-group processes that are unique to social species and have no analog in metapopulations (e.g., fissions, fusions, and between-group competition). Herein lies the primary novelty of our work by considering how processes occurring at the group level itself influences dynamics.

### 4.1 Baseline: Without between-group processes

We begin with only births and deaths within the group (no between-group processes), so groups are entirely independent. Groups will eventually go extinct with probability 1 as a result of demographic stochasticity. Because there are no processes to return lost groups to the population, the result is exponential decay at the population level (Fig. 2a). The rate of exponential decay is influenced by within-group density dependence. When groups grow according to an Allee effect, the population shrinks more rapidly (Fig. 2a, b). As the groups’ carrying capacity increases, the per capita population growth rate (Eq. 22) increases, but always remains negative (Fig. 2b). The effect of within-group density dependence on population persistence can be understood through its influence on the time it takes until each individual group goes extinct due to stochastic fluctuations. Growing logistically and increasing the carrying capacity increases the persistence time of groups (Fig. 2c). Likewise, within-group density dependence has a large influence on the distribution of groups. If the carrying capacity is small enough, an Allee effect can result in smaller groups always being more frequent than groups at the carrying capacity in the population (Fig. 2d). In contrast, logistic growth within groups (or even an Allee effect with a large enough carrying capacity), tends to result in most groups being near their carrying capacity (Fig. 2e).

**Figure 2:**
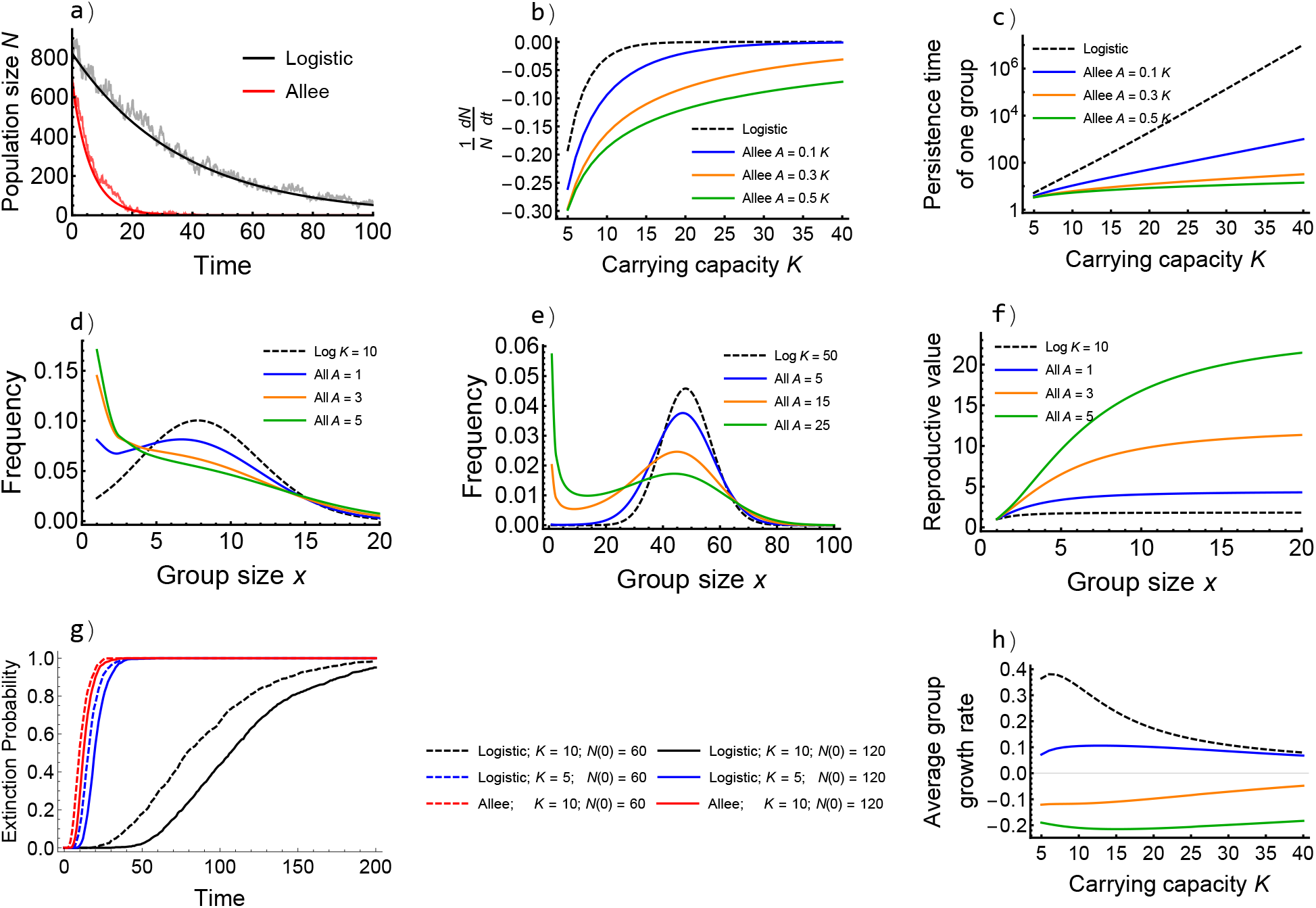
Results from the baseline model without between-group processes. (a) Timeseries showing the agreement between the deterministic (dark) and stochastic (pale) models with logistic (black) and Allee (red) dynamics within the group for *K* = 10 and *A* = 3. (b) Per capita population growth rate increases to 0 as carrying capacity increases (see in-panel legend for form of within-group density dependence). (c) Persistence time of a single group increases exponentially with carrying capacity and is much longer if groups grow logistically. (d) Group size distribution with carrying capacity *K* = 10. (e) Group size distribution with carrying capacity *K* = 50. (f) Reproductive value of a group increases with group size *x*, especially when dynamics are given by an Allee effect. (g) Extinction risk of the entire population is determined more by within-group dynamics than initial population size, *N* (0) (all groups initially at carrying capacity). (h) Average group growth rate increases with carrying capacity *K* given an Allee effect but declines with *K* for logistic growth (coloring same as panels b and c). All panels: *b*_0_ = 3, *d*_0_ = 1.

The simple fact that small groups are prone to extinction has many implications. Large groups have a higher reproductive value because they will always, on average, take longer to go extinct. This is especially true when groups grow according to an Allee effect since small groups are far more extinction prone under these conditions (Fig. 2f). Another effect of heightened extinction risk in small groups is that, at the population level, extinction risk is more strongly influenced by within-group density dependence than initial population size (Fig. 2g). Though increasing the initial population size (by including more groups at carrying capacity initially) decreases extinction probability, the effect is minor (compare solid to dashed lines of each color in Fig. 2g). In contrast, either halving the group carrying capacity (but introducing more groups) or switching within-group dynamics to an Allee effect results in a large increase in extinction probability (compare lines of different colors in Fig. 2g). These results imply that small populations consisting of large groups are far less extinction prone than large populations consisting of small groups. Again, this occurs because small groups (and groups with an Allee effect) persist for much shorter periods of time before extinction due to demographic stochasticity. From the perspective of the individual, however, there may be a benefit to smaller carrying capacities. When groups grow logistically, the average group growth rate is largest in groups with small carrying capacity. In contrast, the average group growth rate of groups with an Allee effect increases slightly with carrying capacity, because there are fewer groups below the Allee threshold with larger carrying capacities (Fig. 2h).

### 4.2 Effect of each between-group process

We now characterize the effect of each between-group process on dynamics, beginning with the well-understood processes of migration and extinction and continuing next to the more novel features of our models. Results are summarized in Table 3. We focus on each process in isolation to allow for a thorough treatment of its role in dynamics.

**Table 3:** Summary of the effect of increasing the rate of each between-group process (relative to the baseline); numbers correspond to section in which relevant analysis described.

| Process | Group size distribution<br>(3.2.2) | Population growth rate<br>(3.2.2) | Average group growth rate<br>(3.2.2) | Group persistence time<br>(3.3.3) | Population extinction risk<br>(3.1.2) |
| --- | --- | --- | --- | --- | --- |
| Migration | Shifts right<br>(more large groups) | Increases | Logistic: Decreases;<br>Allee: Increases | Increases | Decreases |
| Group extinction | No change | Decreases | No change | Decreases | Increases |
| Fission | Shifts left<br>(more small groups) | Increases | Logistic or Allee w/<br>small $A$: Increases;<br>Allee w/ large $A$:<br>Decreases | Decreases | Logistic: Decreases;<br>Allee: Lower w/<br>intermediate fission |
| Fusion | Shifts right<br>transiently | Decreases | Decreases | Increases | Increases |
| Group competition | Shifts right<br>transiently | Decreases | Decreases | Decreases | Increases |

#### 4.2.1 Migration

The overall effect of most forms of migration is to generate a net flux of individuals from large to small groups. This occurs with a constant per capita emigration rate (since there are more individuals to leave from large groups, black line; Fig. 3a) or a per capita emigration rate that increases with group size (blue line; Fig. 3a). If, however, the per capita emigration rate scales with 1/*x* (decreases in large groups), then the rate at which any individual leaves a group is constant and thus the net immigration rate is always 0 (red line; Fig. 3a). When the net immigration rate decreases from positive to negative with increasing group size, small groups become larger and large groups smaller. This causes a contraction in the width of the group size distribution. However, the contraction is not symmetric; small groups grow more than large groups shrink. This is because adding one individual to a small group constitutes a much greater relative change in group size (and thus group growth rate) than removing one individual from a large group. As a result, the group size distribution shifts to the right such that there are fewer small groups in the population (Fig. 3b), an effect that is especially prevalent when groups grow according to an Allee effect (Fig. 3c). The effect of decreasing the number of small groups is to decrease the population’s extinction risk (Fig. 3d). This is referred to as the “rescue effect” in metapopulation ecology (Brown and Kodric-Brown, 1977; Eriksson et al., 2014). It occurs because small groups grow, and thus become less likely to go extinct, when they receive immigrants. This means that the persistence time of any one group increases with migration rate (Fig. 3e).

**Figure 3:**
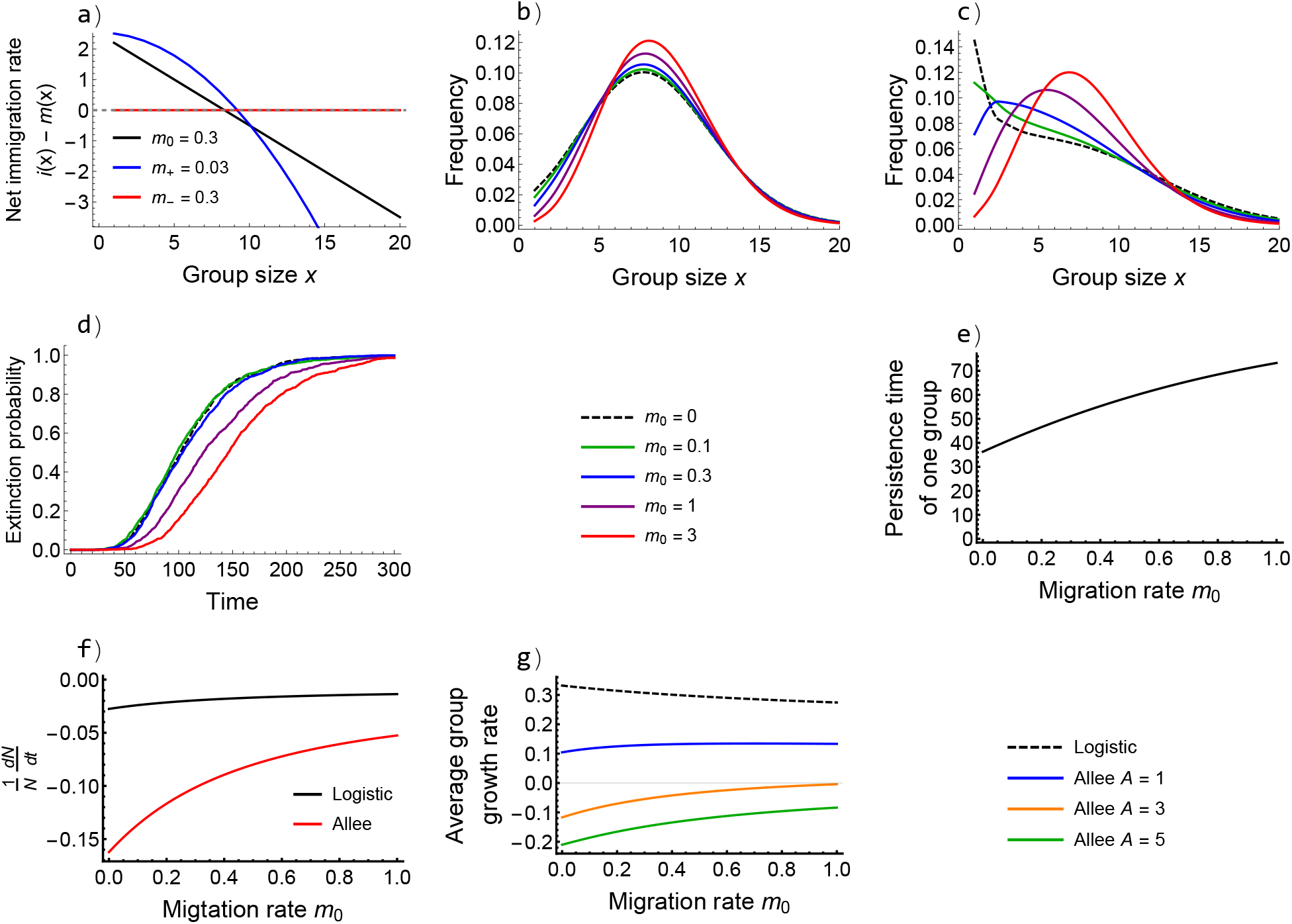
The effect of migration on population dynamics. (a) The net immigration rate (*i*(*x*) *− m*(*x*)) across group sizes. When the per capita migration rate is constant (black line) or increases with group size (blue line), migration causes a net flux of individuals from large to small groups. If the per capita migration rate is highest in small groups (red line), then the net migration rate is 0 for all group sizes. (b) Higher migration rates shift the group size distribution to the right with logistic growth within groups (c) and especially so with an Allee effect within groups. Increasing migration rates also (d) decrease the population’s extinction risk, (e) increase the time for which a single group persists, and (f) increase the per capita population growth rate. (g) The effect of migration on the average group growth rate depends on within-group density dependence. All panels: *b*_0_ = 3, *d*_0_ = 1, *K* = 10, with logistic growth (unless otherwise specified, all unspecified parameters equal to 0).

By shifting the group size distribution to the right and leading to fewer small groups, migration has predictable effects on growth at both the group and population level. Because groups go extinct more slowly, increasing the migration rate increases the per capita population growth rate (Fig. 3f). Whether or not the average group growth rate increases or decreases depends upon within-group density dependence. With logistic growth, increasing migration rates cause a slight decrease in the average group growth rate, because of stronger negative density dependence in larger groups (black line; Fig. 3g). In contrast, with a high Allee threshold, increasing the migration rate considerably decreases the proportion of groups that are trapped below the Allee threshold, thus increasing the average group growth rate (blue, orange, and green lines; Fig. 3g).

#### 4.2.2 Extinction

Unsurprisingly, increasing extinction rates mean that groups are removed from the population more rapidly, so each group persists for considerably less time (Fig. 4a). As we have seen previously, group extinction is the source of permanent population decline in social species and thus this decreases the per capita population growth rate (Fig. 4b). Extinctions are less detrimental when they occur less frequently from large groups, as small groups are likely to go extinct quickly anyway due to demographic stochasticity (compare red and black lines in Fig. 4a,b). If extinctions occur independent of group size they have no effect on the group size distribution (Fig. 4c). In contrast, if small groups are more likely to go extinct this will shift the group size distribution to the right, but the effect is relatively small unless the extinction rate is extremely high (Fig. 4d).

**Figure 4:**
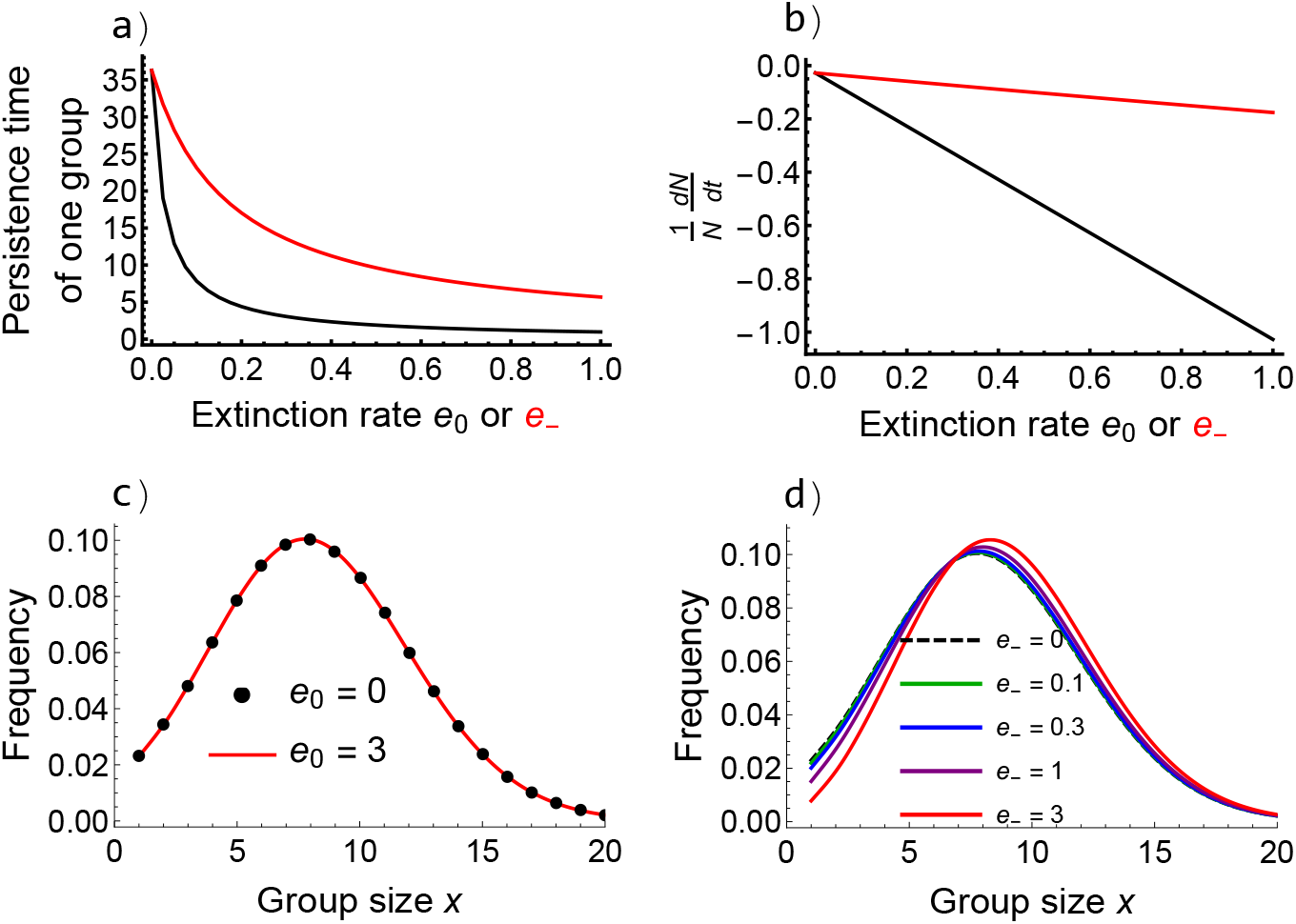
The effect of group extinctions on population dynamics. Increasing extinction rate (a) decreases the persistence time of groups and (b) decreases the per capita population growth rate. (c) When extinctions occur independent of group size, the group size distribution does not change, (d) but when extinction of small groups is more likely the group size distribution shifts slightly to the right. All panels: *b*_0_ = 3, *d*_0_ = 1, *K* = 10, logistic growth, unspecified parameters equal to 0.

#### 4.2.3 Fission

The most obvious effect of fissions is that they generate new groups. If a single group is expected to fission at least once before going extinct, then the number of groups in the population will increase exponentially (blue line; Fig. 5a). In addition to increasing numbers of groups, fissions decrease the average group size (red line; Fig. 5a) by generating an excess of small groups (Fig. 5b). Since intragroup competition is alleviated in these small groups, they are expected to grow and return to their carrying capacity. Thus, with fissions, the total population size can increase exponentially as well (black line; Fig. 5a). Consequently, fissions increase the per capita population growth rate (Fig. 5c). This qualitative effect of fissions occurs with or without group size heterogeneity, but is artificially strong if we ignore heterogeneity since doing so spreads the density-dependent benefits of smaller post-fission group sizes across all groups. With an Allee effect, we see some qualitative influence of group size heterogeneity. When the fission rate becomes too high population growth does not keep up with group growth without explicitly modeling multiple groups and thus the population crashes once fission rates become too high (dashed red line; Fig. 5c). Regardless of group size heterogeneity, density dependence is lacking from the population level with fissions since the rate (and effect) of a single fission does not depend upon the population size. With fissions, large groups still contribute more to population growth (have higher reproductive value), but this effect is weaker than when fissions are absent (reproductive value increases more slowly with group size; Fig. 5d). The reason is that reproductive value is determined both by the generation of new groups and persistence. Group fissions provide a means for all groups of more than 1 individual to generate new groups, and make it so that large groups are unlikely to remain large for long.

**Figure 5:**
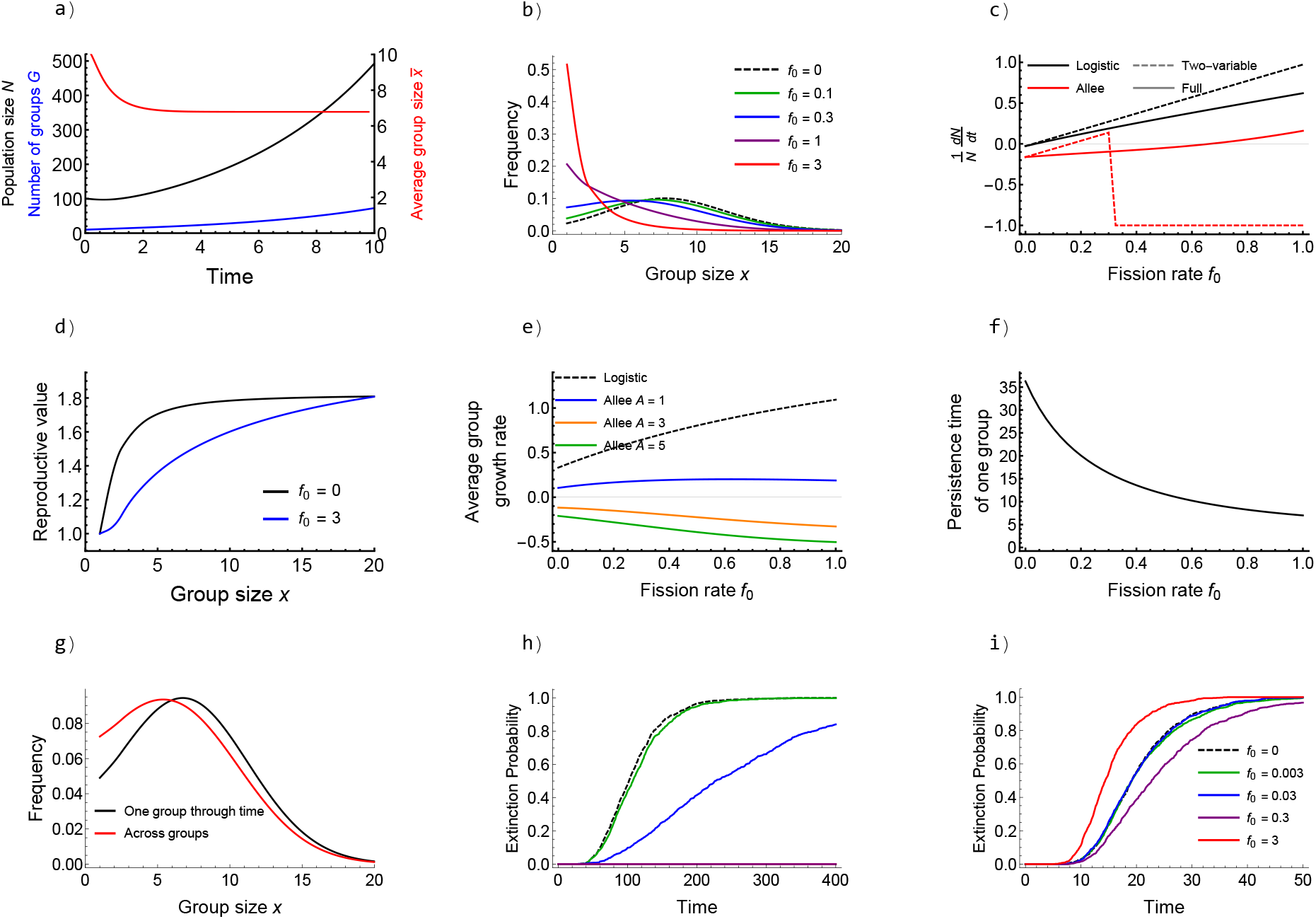
The effect of fissions on population dynamics. (a) Timeseries with fissions at rate *f*_0_ = 0.3 showing that group number (blue line) and population size (black line) increase with time and average group size (red line) decreases to below the carrying capacity *K* = 10. (b) Higher fission rates shift the group size distribution to have more small groups. (c) The per capita population growth rate increases with fission rate. (d) When fissions occur, increasing reproductive value with increasing group size is slower than without fissions. (e) The effect of fission rate on average group growth rate depends on within-group density dependence. (f) Higher fission rates decrease the persistence time of groups. (g) The distribution of a single group through time (black line) is shifted to the right of the population-level group size distribution (red line). (h) Higher fission rates decrease the extinction probability of populations with logistically growing groups (see i for legend) and (i) initially with groups that have an Allee effect. But with an Allee effect and high enough fission rates, fissions increase extinction probability. All panels: *b*_0_ = 3, *d*_0_ = 1, *K* = 10, *A* = 3, logistic growth (unless otherwise specified), unspecified parameters equal to 0. Qualitative conclusions remain when fission rates depend on group size (i.e., *f*_+_ > 0).

Despite these clear benefits to population growth, within groups the effect of fissions is more nuanced and mediated by within-group density dependence. The result of making the average group size smaller is that groups spend more time growing if within-group dynamics follow logistic growth (the average group growth rate increases; Fig. 5e). However, if there exists an Allee threshold *A*, high fission rates often cause groups to be smaller than *A*, leading to a decrease of average group growth rate with fission rate (Fig. 5e). By generating more small groups, fissions decrease the expected time until any one group goes extinct (Fig. 5f). The shorter persistence of small groups also means that the average group spends most of its lifetime at larger sizes; if it becomes small (e.g. as a result of fission), it is unlikely to remain small for long before either going extinct or escaping extinction by growing large again. So, the distribution of any individual group’s size throughout its lifetime is shifted to the right relative to the distribution of group sizes in the population at any one moment in time (i.e., the model is no longer ergodic; Fig. 5g). Put another way, although small post-fission groups are well represented in the distribution of group sizes in the population at any given moment, their short longevity diminishes their impact on the temporal distribution of sizes experienced by any individual group.

The dual nature of small groups also emerges to influence extinction risk at the population level. If groups grow logistically, the increase in the number of groups as a result of increasing fission results in populations being less extinction prone (Fig. 5h). When groups grow according to an Allee effect, likewise, increasing fission rate can provide a means to “replace” extinct groups for sufficiently low fission rates, decreasing extinction probability (purple line; Fig. 5i). However, if the fission rate becomes too high, too many groups are created that are below their Allee threshold and each group becomes much more likely to go extinct before returning to carrying capacity, thus driving an increased extinction risk when fissions occur at a sufficiently high rate (red line; Fig. 5i).

#### 4.2.4 Fusion

The influence of fusions depends upon the total number of groups in the population (*r* and *ψ* change with *θ* not just 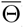). This means that fusions are both density dependent (their influence on population growth rate changes with population size) and that their effects are transient in exponentially declining populations (Fig. 6a, because fusions become increasingly rare as the population density goes to 0). In contrast with fissions, fusions create fewer (and larger) groups in the population (Fig. 6a). Because density dependence occurs within groups, fewer groups support fewer individuals at the population level and thus increasing fusion rate decreases the population growth rate (black line Fig. 6b); a result that does not depend upon heterogeneity in group size (dashed line Fig. 6b). Since population extinction relies mainly upon the number of groups (Fig. 2g), increasing the fusion rate also increases the population’s extinction risk (Fig. 6c). If, however, fusions are only likely to occur with small groups (*ρ*_−_ > *ρ*_0_ = 0), then the decline in population growth rate observed when larger groups fuse is mitigated since small groups are likely to go extinct anyway (red line Fig. 6b). Still, this does not permit a benefit to population growth over the no-fusion scenario because a fusion results in two groups (stable at 2*K* individuals) becoming one (stable at *K* individuals) and thus is no different in the long run than the extinction of one of the groups: any benefit is inherently transient.

**Figure 6:**
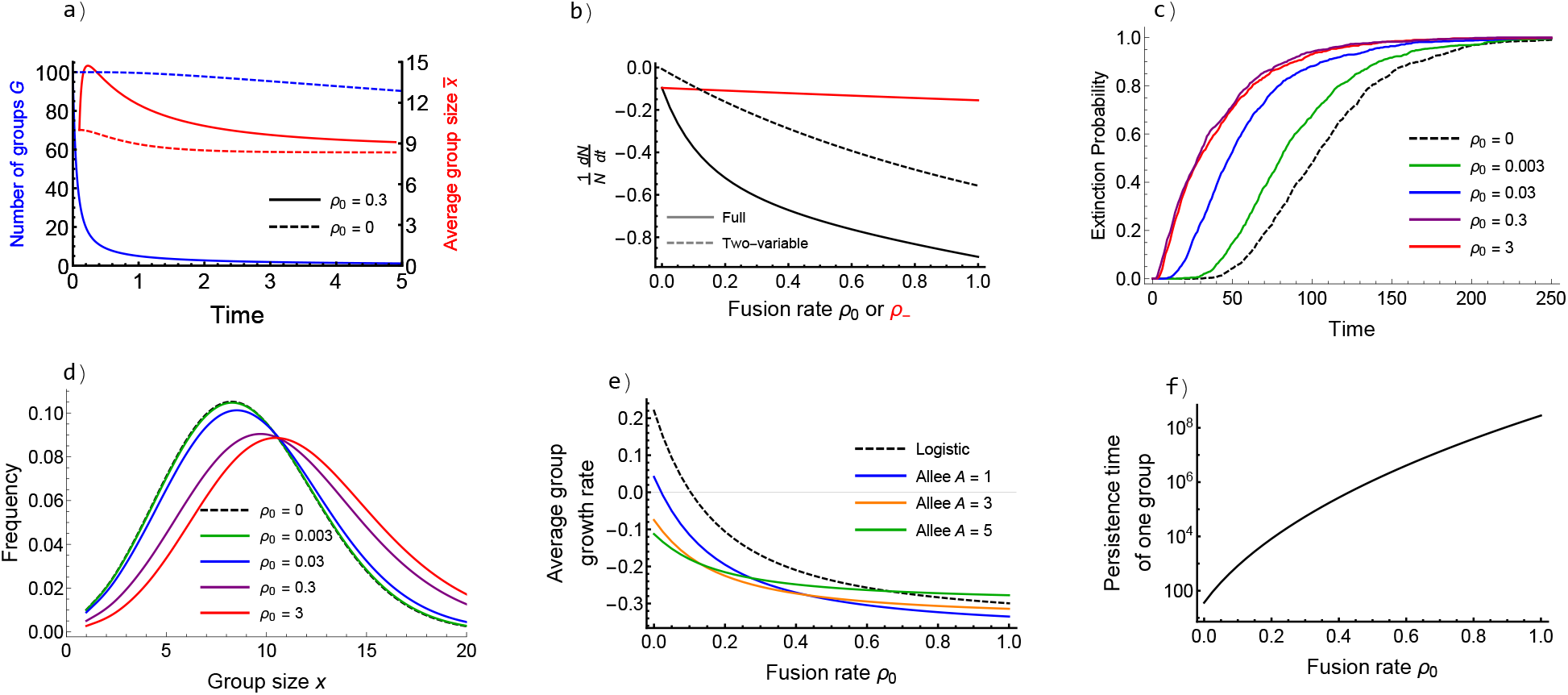
The effect of fusions on population dynamics. (a) Timeseries contrasting a population without fusions (dashed) and a population with fusions (solid). Fusions cause the number of groups to rapidly decrease (blue) increasing the average group size above carrying capacity *K* = 10 though the effect is transient (solid red line approaches dashed red line). (b) Higher fusion rates that are independent of group size transiently decrease the per capita population growth rate (black; shown here after 1 time unit such that each group of size *x* is expected to fuse once when *r*(*x*) = 1/*G*), but when fusions only occur in small groups the effect is negligible (red). Qualitative results do not depend on group size heterogeneity (see dashed line from the two-variable model). Higher fusion rates (c) lead to heightened extinction risk, (d) transiently shift the group size distribution to the right (shown after 1 time unit), (e) decrease the average group growth rate (shown after 1 time unit), and (f) increase the time for which a single group persists. All panels: *b*_0_ = 3, *d*_0_ = 1, *K* = 10, logistic growth (unless otherwise specified), unspecified parameters equal to 0.

By creating larger groups upon a fusion, increasing the fusion rate (transiently) shifts the group size distribution to the right, potentially exceeding the within-group carrying capacity (Fig. 6d). The result is that the average growth rate of groups also decreases with increasing fusion rate (Fig. 6e). Shifting the group size distribution to the right does provide some benefit as it increases the persistence time of groups (Fig. 6f). As explained above, increasing the extinction time of groups through fusions does not considerably change dynamics because fusions decrease the total potential number of individuals that can stably exist in the population as a result of group-level negative density dependence.

#### 4.2.5 Group competition

As with fusions, the effects of group competition are transient in exponentially declining populations as the interaction rate between groups goes to 0 as the number of groups in the population goes to 0. Depending on the effect of group competition on groups, its effect on the group size distribution changes. If competition results in a single death (our main modeling assumption), then small groups become over-represented (Fig. 7a). However, if competition results in the extinction of an entire group, then large groups become over-represented (Fig. 7b). Group competition has a much larger impact on the group size distribution when only a single individual dies because in this case it fundamentally alters the realized within-group density dependence. Though other results between these model variants do not differ qualitatively (and thus we show only results for when competition results in a single death), this demonstrates that the way competition alters the group size distribution in real systems is contingent upon the form of competition. As expected, because it is a source of additional mortality in the population, group competition decreases the per capita population growth rate (Fig. 7c). Importantly, however, this effect is now density dependent at the population level, unlike all other sources of mortality considered. Because the rate at which groups interact increases with the number of groups in the population, the added death rate from group competition increases with population size (Fig. 7c).

**Figure 7:**
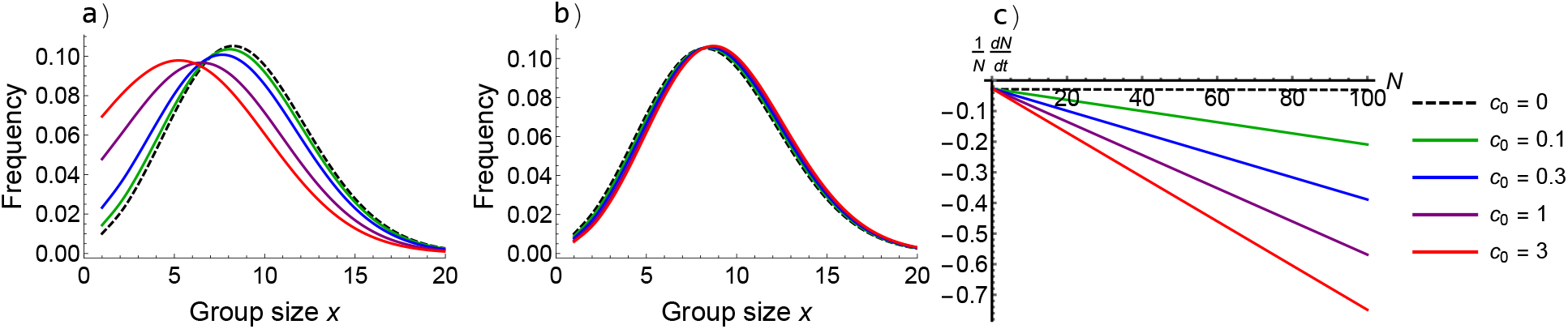
The effect of group competition on population dynamics. The effect of group competition on the group size distribution depends on the way that competition influences the population. (a) When competition results in a single death, small groups are over-represented. (b) When competition results in group extinction, large groups are over-represented. (c) Group competition leads to density dependence at the population level, with higher competition rates resulting in lower per capita population growth rates. All panels: *b*_0_ = 3, *d*_0_ = 1, *K* = 10, logistic growth (though qualitative results hold with an Allee effect), unspecified parameters equal to 0. (a) and (b) use *t* = 1. (c) uses *t* = 100.

### 4.3 Interactions of between-group processes

Now that we have analyzed the effect of each of the five between-group processes on dynamics, we turn to understanding interactions among these processes. Unlike above, we do not attempt a thorough investigation of each possible interaction as this quickly becomes unruly (there are 10 possible pairwise interactions alone). Rather, we focus on two special cases that are likely present in nature and can be expected to have important impacts on dynamics: interactions between fissions and fusions as well as fissions and group competition.

#### 4.3.1 Fission and fusions

The fact that fusions occur at a greater rate when there are more groups in the population (are density dependent at the population level) is an important mediator of dynamics. Populations with groups that both fission and fuse display logistic growth at the population level (Fig. 8a-d), even if groups grow according to an Allee effect (Fig. 8b,d). Here, the nature of between-group processes leads to population regulation without there being any explicit form of competition that depends upon the total number of individuals in the population. This occurs whenever fission rates are high enough that populations would grow exponentially in the absence of other between-group processes. In this case, fusions are rare in small populations and thus unlikely to occur, meaning that the number of groups will increase in small populations since the total rate of fissions is greater than the total rate of fusions. Spreading the same number of individuals into more groups also increases the per capita population growth rate due to weaker intragroup competition and within-group density dependence. However, the rate of fissions increases linearly with group size, whereas the rate of fusions increases quadratically with group size. Consequently, once the population size is large enough, the rate of fusions will be greater than the rate of fissions. Once this population size is reached, the rate of change of the number of groups, and thus the population growth rate, becomes negative (Fig. 8c,d). This demonstrates that population-level density dependence in social species can emerge due to the dynamics of social groups themselves. Here, given the existence of processes that limit group size, patterns of births and deaths occurring within groups have no influence on the form of density dependence that emerges at the population level.

**Figure 8:**
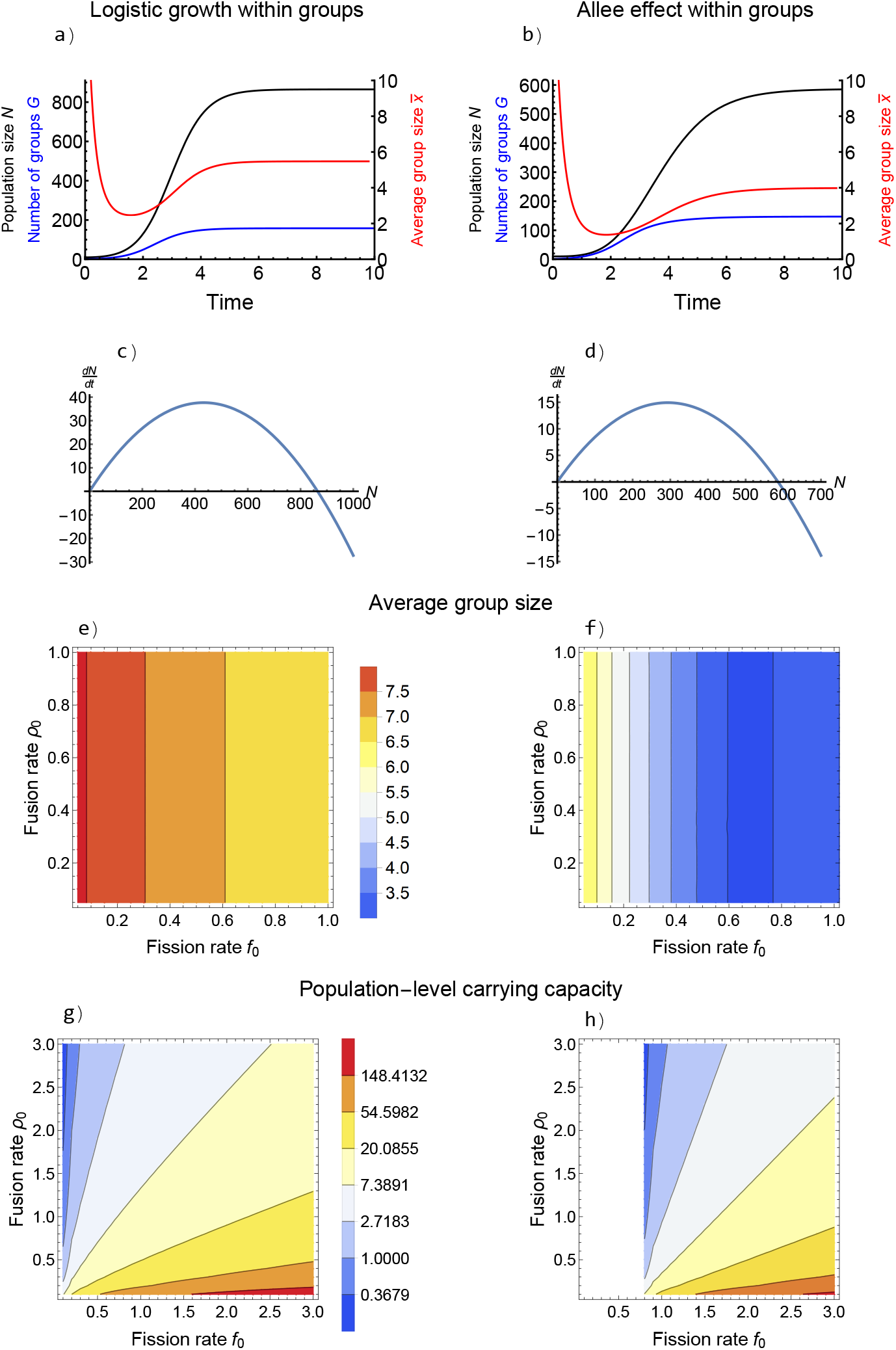
Interactions between fissions and fusions. Throughout, the left-hand column shows logistic growth within groups with *b*_0_ = 3, *d*_0_ = 1, *K* = 10 and the right-hand column shows an Allee effect within groups with *b*_0_ = 3, *d*_0_ = 1, *K* = 10, *A* = 3. (a, b) Timeseries demonstrating logistic growth at the population level (*f*_0_ = 3 and *ρ*_0_ = 0.03). (b, c) Population growth rate versus population size (assuming constant group size distribution; same parameters as above). (e, f) The average group size at equilibrium (red is larger; see legend) decreases with fission rate and is smaller when groups grow according to an Allee effect. (g, h) Higher fission rates and lower fusion rates lead to a higher population-level carrying capacity (red is larger; see legend). White regions correspond to parameter combinations that result in population extinction.

Though the qualitative dynamics of fissions and fusions at the population level hold across parameter values and with either form of within-group density density dependence, these features do influence both the average group size at equilibrium and the value of the population-level carrying capacity. First, average group size at equilibrium depends upon the fission rate, but not the fusion rate, and decreases with increasing fission rates (Fig. 8e,f). The lack of dependence on the fusion parameter *ρ*_0_ arises because the realized fusion rate experienced by any one group is not actually *ρ*_0_ but rather *ρ*_0_*G*. The number of groups *G* at equilibrium decreases with increasing fusion rates and this offsets to keep *ρ*_0_*G* nearly constant, meaning that the fusion rate parameter itself has little influence on the equilibrium average group size. The average group size at equilibrium is also substantially smaller when groups grow according to an Allee effect (Fig. 8e,f), since high fission rates will result in many groups below the Allee threshold. Second, the population-level carrying capacity increases with fission rate and decreases with fusion rate (Fig. 8g,h). This can be understood by the metaphor that fissions are like the birth of new groups and fusions like the death of groups. The population-level carrying capacity is slightly smaller when groups grow according to an Allee effect compared to logistically for a given set of parameters, but the effect is small (Fig. 8g,h). Group size heterogeneity is not required for fissions and fusions to result in logistic growth at the population level. The two-variable model can be treated analytically in this case and, assuming logistic growth, has equilibrium 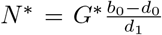 and 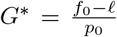. The only qualitative difference is that spreading the effect of between-group processes over all groups in the two-variable model means they have no effect on the average group size at equilibrium. Taken together, these results demonstrate that population-level density dependence is decoupled from within-group dynamics.

#### 4.3.2 Fissions and group competition

Interactions between fissions and group competition are qualitatively the same as those between fissions and fusions. Most notably, if the fission rate is high enough to permit exponential growth without other group-level processes, then again logistic growth will emerge at the population level when we add group competition (Fig. 9a,b). Now, this occurs because the death rate increases as the number of groups in the population increases (due to intergroup competitions), which in turn increases the rate at which groups are lost from the population. Notably, this feedback does not occur in the two-variable model without group size heterogeneity because increasing the death rate (the effect of group competition) does not increase the rate at which groups are lost in this model, and thus logistic growth does not emerge at the population level with fissions and group competition here. Again, the fission rate is the primary determinant of the average group size at equilibrium, which decreases with increasing fission rate. There is a small effect of increasing competition rates decreasing the average group size at equilibrium that only occurs when the competition rate is low (Fig. 9c). The population-level carrying capacity increases with increasing fission rate and decreases with increasing competition rate. It appears most sensitive to the competition rate, only increasing with the fission rate when the fission rate is relatively low (Fig. 9d).

**Figure 9:**
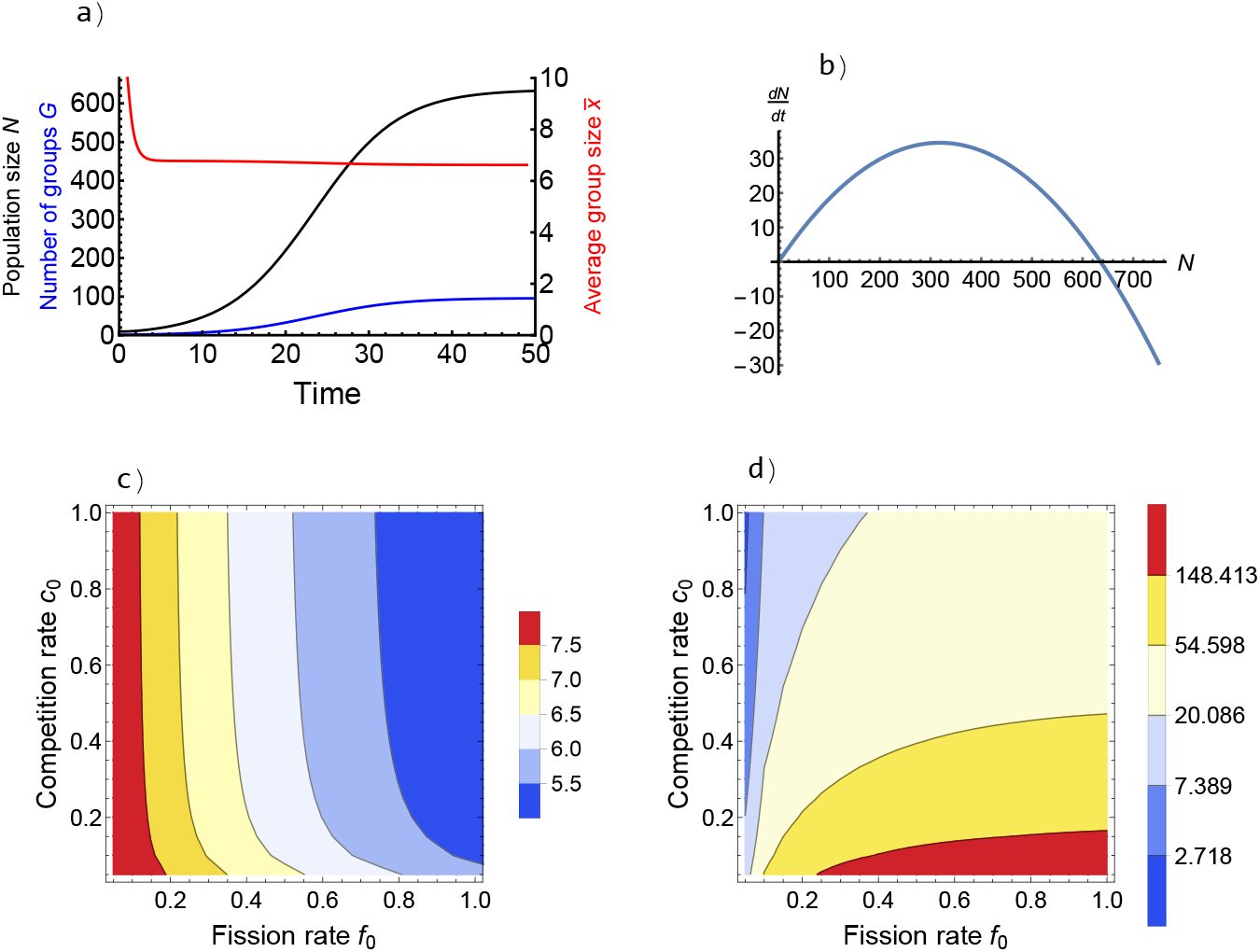
Interactions between fissions and group competition in logistically growing groups (dynamics are similar for groups with an Allee effect). (a) Timeseries showing logistic growth at the population level. (b) Population growth rate versus population size. (c) Average group size at equilibrium (red is larger; see legend) depends mostly on fission rate and decreases with fission rate. (d) Population-level carrying capacity (red is larger; see legend) increases with fission rate (especially when the fission rate is low) and decreases with competition rate. *b*_0_ = 3, *d*_0_ = 1, *K* = 10.

## 5 Modeling distinct ecological roles

So far, we have seen how patterns of within-group density dependence and between-group processes shape the dynamics of social populations. Throughout, we have made the simplifying assumption that all individuals are ecologically identical. Missing from the previous results is the idea that individuals play distinct ecological roles within their group (e.g., due to age, sex, or class structure), which, in many social animals, are already known to be key drivers of dynamics. Only by permitting distinct ecological roles can our framework capture the importance of offspring provisioning (a driver of cooperative breeding; Lukas and Clutton-Brock, 2012), the dependence of new group formation on juvenile dispersal seen in some species (McNutt, 1996), ecologically relevant sex roles in primates (Kappeler, 2017), the ecological implications of the common pattern of sex-biased dispersal (Trochet et al., 2016), and the division of labor that makes eusocial animals unique (Crespi and Yanega, 1995). Accounting for such individual differences thus allows us to create a truly general mathematical tool for studying social populations: a primary goal of this study.

### 5.1 Overview

In the evolutionary models that motivated our own (Simon, 2010; Simon et al., 2013), multiple types of individuals were included as different phenotypic strategies. Here, following this previous work, we show how this framework can be extended to consider an arbitrary number of types (*x*_1_, *x*_2_,…), in our case different classes of individuals. As in section 3.1, we begin with a stochastic version of the model with distinct ecological roles that can be simulated with Gillespie’s stochastic simulation algorithm. Now, we track the number of groups with *x*_1_ individuals of type 1, *x*_2_ individuals of type 2, and so on. Again, taking the number of groups to infinity appropriately leads to the Markov chain converging to a system of ODEs. The number of state variables that must be tracked is 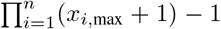, where *n* is the number of types and *x_i,_*_max_ is the maximum number of individuals of type *i* that can be present in a single group. The number of states clearly grows rapidly with *x_i,_*_max_ and *n*. Now, it is possible to enter the state (*x*_1_, *x*_2_, …, *x*_*n*_) by changing the number of type 1 individuals in the group, type 2 individuals in the group, and so on (including changing multiple types of individuals within the group). It is again possible to obtain results either by working with a rate matrix in linear cases or by numerically integrating the system of ODEs in all cases.

We now show the implications of individuals of social species having distinct ecological roles with a series of empirically-relevant case studies that consider age-structured groups, sex-structured groups, and class-structured groups. Our goal is not necessarily to provide insight into any real system, but rather to demonstrate the flexibility of this approach. For tractability, we restrict ourselves to groups containing two types of individuals. We begin each case study with a brief overview of modeling assumptions before showing results.

### 5.2 Age-structured groups

We first consider two case studies for the role of age structure in social populations. We will refer to juveniles (individuals that are not yet sexually mature) as type 1 and adults (sexually mature individuals) as type 2.

#### 5.2.1 Case study I: Age structure and growth rates

To begin, we present a model that is as similar as possible to our baseline case (except for the addition of age structure) to provide the best comparison for how age structure affects our results. We assume that each adult gives birth to a juvenile at a per capita rate of *b*_0_. Adults die according to the same per capita death rate as logistic growth from our model with ecologically identical individuals, with density dependence imposed only by other adults, such that

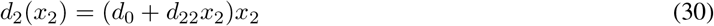

is the overall rate at which adults die in a group with *x*_2_ adults. We assume, on the other hand, that more adults in the group is beneficial for juveniles such that the juvenile death rate is

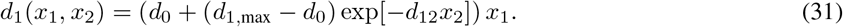

Here, the per capita juvenile death rate is a decreasing function of the number of adults in the group (approaching the minimum adult death rate *d*_0_ in the limit of infinite adults) with a maximum of *d*_1,max_ when there are no adults. Finally, we assume that juveniles mature into adults at a constant per capita rate *μ*.

When the maturation rate is slower than the birth rate, including age structure reduces the per capita population growth rate until it approaches the per capita population growth rate from the model without age structure as maturation becomes very fast (Fig. 10a). The persistence time of groups follows a similar pattern, with the longest persistence times following from high maturation rates (Fig. 10c). At low maturation rates, maturation is not rapid enough to replenish adult deaths (Fig. 10d), and thus groups often go extinct (Fig. 10c) with the population shrinking rapidly (Fig. 10g). At high maturation rates, individuals spend very little time is spent as juveniles. Consequently, groups consist of few juveniles (Fig. 10f) and age structure has little influence on dynamics, so results converge to those seen in the case without age structure (Fig. 10a-c). As a result, the per capita population growth rate become almost independent of juvenile density dependence (Fig. 10i). When the maturation rate is comparable to the birth rate, groups consist of approximately equal numbers of juveniles and adults (Fig. 10e). Groups with more adults have higher birth rates and juveniles in such groups do not suffer from higher death rates, so a positive correlation arises between the number of juveniles in a group and the number of adults in a group in this regime. As a result, average group growth rate is greatest, typically greater than it would be without age structure, when the maturation rate is intermediate (comparable to the birth rate) (Fig. 10b). This occurs because only at intermediate maturation rates do groups consist of a substantial number of both adults and juveniles, which allows for positive density dependence in juvenile mortality to buffer negative density dependence in adult mortality.

**Figure 10:**
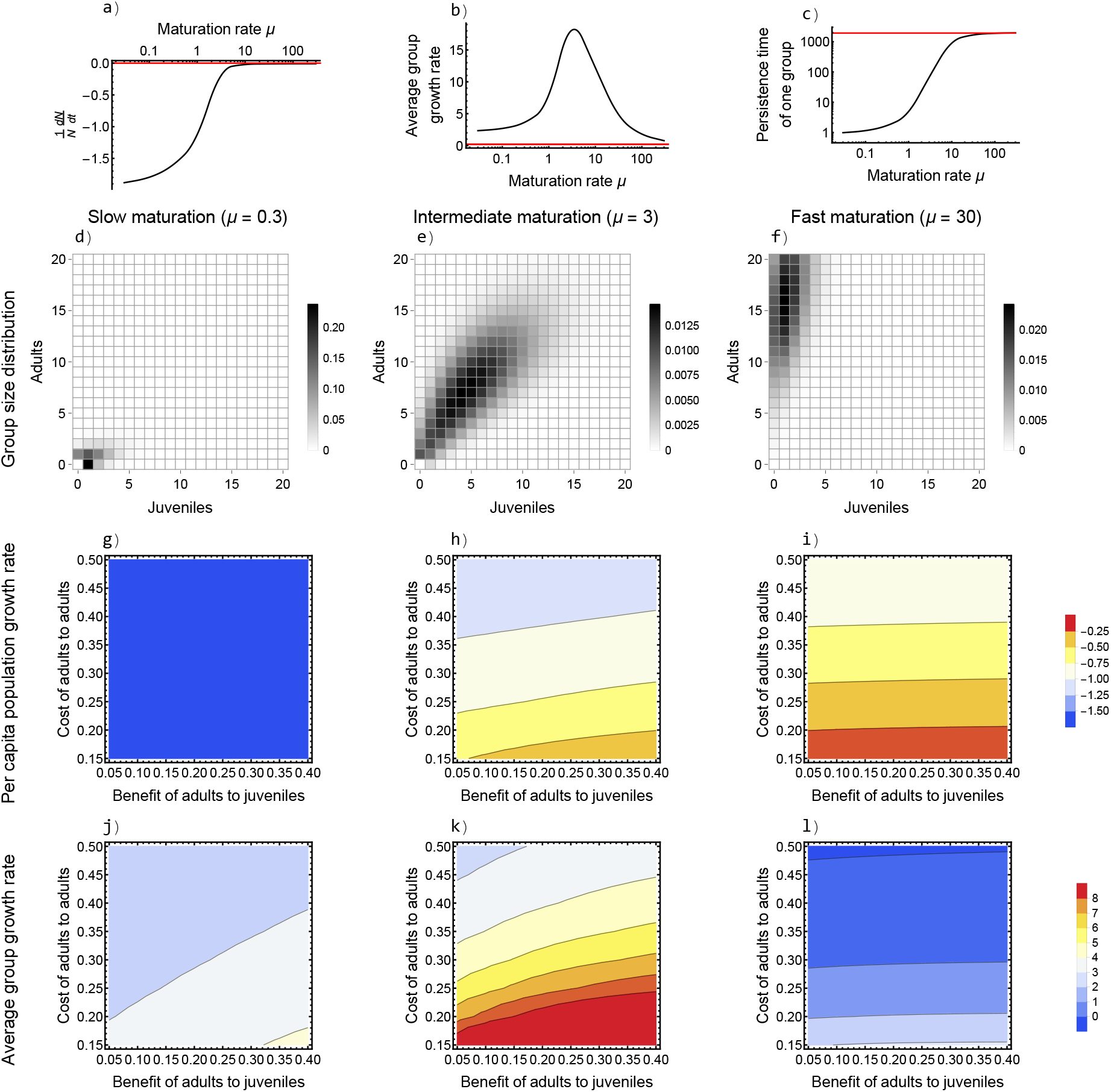
Results from the model with age structure and a density-dependent tradeoff between juvenile and adult mortality. In the first row (a-c), the black lines show results from the model with age structure, while red lines indicate values from the baseline model without age structure with the same parameter values for comparison. (a) Per capita population growth rate increases with increasing maturation rate, saturating at the value from the model without age structure. (b) The average group growth rate tends to be larger with age structure, with an intermediate peak when the maturation rate is approximately the birth rate. (c) The persistence time of groups increases with increasing maturation rate, saturating at the value from the model without age structure. After the first row, the first column (d, g, j) shows the effect of low maturation rates (*μ* = 0.3 = *b*_0_/10), the second column (e, h, k) shows the effect of intermediate maturation rates (*μ* = 3 = *b*_0_), and the third column (f, i, l) shows the effect of high maturation rates (*μ* = 30 = 10*b*_0_). (d-f) The stable group size distribution, with darker colors indicating more groups of that composition. (g-i) Per capita population growth rate with the benefit to juveniles of adults *d*_12_ on the horizontal axis and the cost to adults of increasing adult density *d*_22_ on the vertical axis. Highest growth rates are indicated by redder colors. (j-l) Average group growth rate with the benefit to juveniles of adults *d*_12_ on the horizontal axis and the cost to adults of increasing adult density *d*_22_ on the vertical axis. Highest growth rates again indicated by redder colors. Unless otherwise specified: *b*_0_ = 3, *d*_0_ = 1, *d*_1,max_ = 2, *d*_12_ = 0.1, *d*_22_ = 0.1.

#### 5.2.2 Case study II: Juvenile migration and new group formation

Our next case study shows how one can mechanistically model group formation and is motivated by social carnivorans, in particular African wild dogs. In nature, new groups of wild dogs form by dispersing juvenile cohorts from two different groups fusing (McNutt, 1996; Somers et al., 2008). Here we assume the birth rate saturates with the number of adults in the group such that

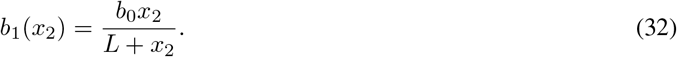

Now, *b*_0_ is the maximum birth rate a group can achieve and *L* is the number of adults such that the birth rate is *b*_0_/2. Death rates follow the same form as Eqns. 30 and 31. The key difference in this model is maturation. We assume that “fissions” occur at rate *f*_+_*x*_1_ and that fissions result in the group’s juveniles and adults separating into one group of only juveniles and one group of only adults. The juvenile-only group only matures (and thus can have births) once it fuses with another juvenile-only group (which occurs at rate 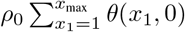). Specifically, we assume that the rate at which a group fuses is 0 if there are any adults in the group (*x*_2_ > 0), and that the fusion of two groups with *u* and *v* juveniles, respectively, results in a single group with *u* + *v* adults.

We find that new group formation via juvenile dispersal requires both sufficiently high fission rates and fusion rates in order for populations to grow (Fig. 11a). The fact that fusions now increase growth corresponds to the fact that new groups cannot reproduce until they fuse with another group. Interestingly, this pattern of new group formation generates positive density dependence at the population level (Fig. 11b; a population-level demographic Allee effect). If too few groups are in the population, then realized fusion rates will never be sufficiently high to permit new groups to begin reproducing due to the lack of juvenile dispersing cohorts. This mirrors the well-known “mate-finding Allee effect” (Gascoigne et al., 2009), by occurring through the inability to find dispersing cohorts when the population is at low density. Such a population-level demographic Allee effect has been shown in a model previously (Courchamp et al., 2000), but was attributed to the scaling of a group-level demographic Allee effect to the population level and stochastic extinction events. Here, we show that a population-level demographic Allee effect can result due to social group dynamics themselves, even with negative density dependence within social groups.

**Figure 11:**
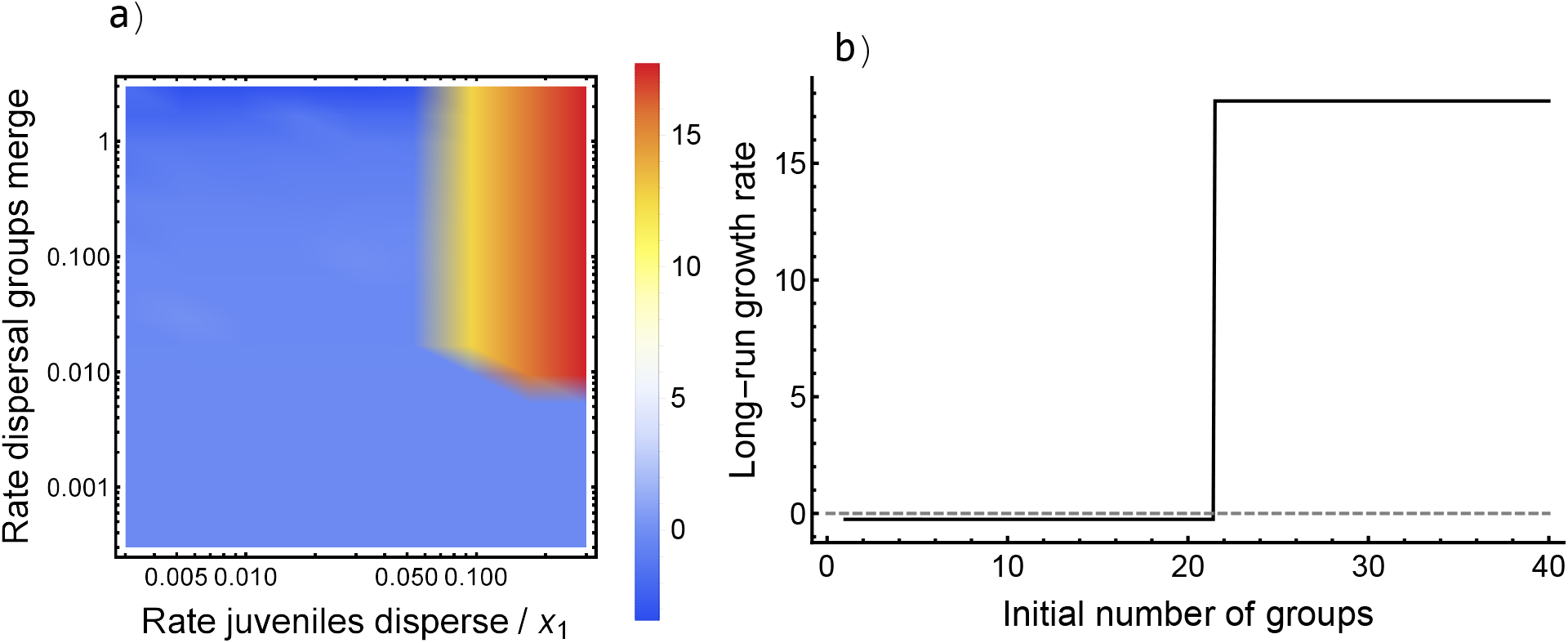
Results from the model with group formation through juvenile dispersal. Positive population growth requires sufficient (a) fission and fusion rates (color shows per capita population growth rate; see legend) and (b) initial number of groups. If the initial number of groups is below a threshold value (here, approximately 21), then the long-run growth rate is negative (extinction); whereas, if there are enough groups initially, then sustained population growth occurs, demonstrating a population-level demographic Allee effect. Unless otherwise specified: *b*_0_ = 50, *L* = 5, *d*_1,max_ = 0.25, *d*_1_2 = 0.25, *d*_0_ = 1, *d*_2_2 = 0.01, *f*_+_ = 0.3, *ρ*_0_ = 0.003.

### 5.3 Sex-structured groups

Our next two case studies consider the implications of two sexes for social populations. We will refer to type 1 individuals as males and type 2 individuals as females.

#### 5.3.1 Case study III: Sex structure and growth rates

The key assumption for sex-structured groups is that the group’s birth rate is determined by the number of females in the group, so long as there is at least one male. That is, the birth rate of type *i* individuals is

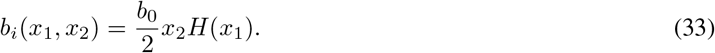

Here, *b*_0_ is the total rate at which a female in a group with at least one male gives birth and the factor of 1/2 accounts for half of births being male and half being female. *H*(*u*) is the Heaviside step function and evaluates to 0 if *u* ≤ 0 and evaluates to 1 otherwise. This accounts for the fact that births can only occur if there is at least one male in the group, but otherwise are independent of the number of males. Deaths for both sexes are assumed to follow logistic growth with an equal competitive effect of both males and females (though potentially different density-independent death rates), such that

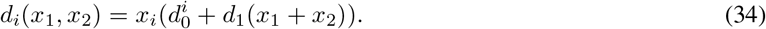

Since males do not contribute to increasing the birth rate (so long as at least one is present), the investment into males resulting from sex structure tends to reduce the population growth rate (related to the “two-fold cost” of the evolution of sex; Maynard Smith, 1971). The group size distribution is skewed, demonstrating that groups with fewer females than males are especially likely to shrink and more females than males especially likely to grow (Fig. 12a; note that groups with more males than females (below the red line) are smaller overall than groups with more females than males (above the red line)). That is, females are more beneficial than males for group growth. One manifestation of this is that groups are more likely to become “trapped” with no females but one or two males (gray squares, bottom left of Fig. 12a), compared to having no males in a group, since groups with few females see a decline in birth rate that groups with few males do not. These results stem from males contributing to competition within the group without contributing to the birth rate. For this reason, increasing the minimum male death rate *d*^1^, may actually increase the per capita population growth rate, because this skews the sex ratio to be female-biased which allows for more births within the group (Fig. 12b). Increasing the minimum female death rate *d*^2^, on the other hand, substantially decreases the per capita population growth rate (Fig. 12b). Another manifestation of the birth rate being determined primarily by the number of females in the group is that reproductive value declines more rapidly in groups with fewer females than fewer males (Fig. 12c).

**Figure 12:**
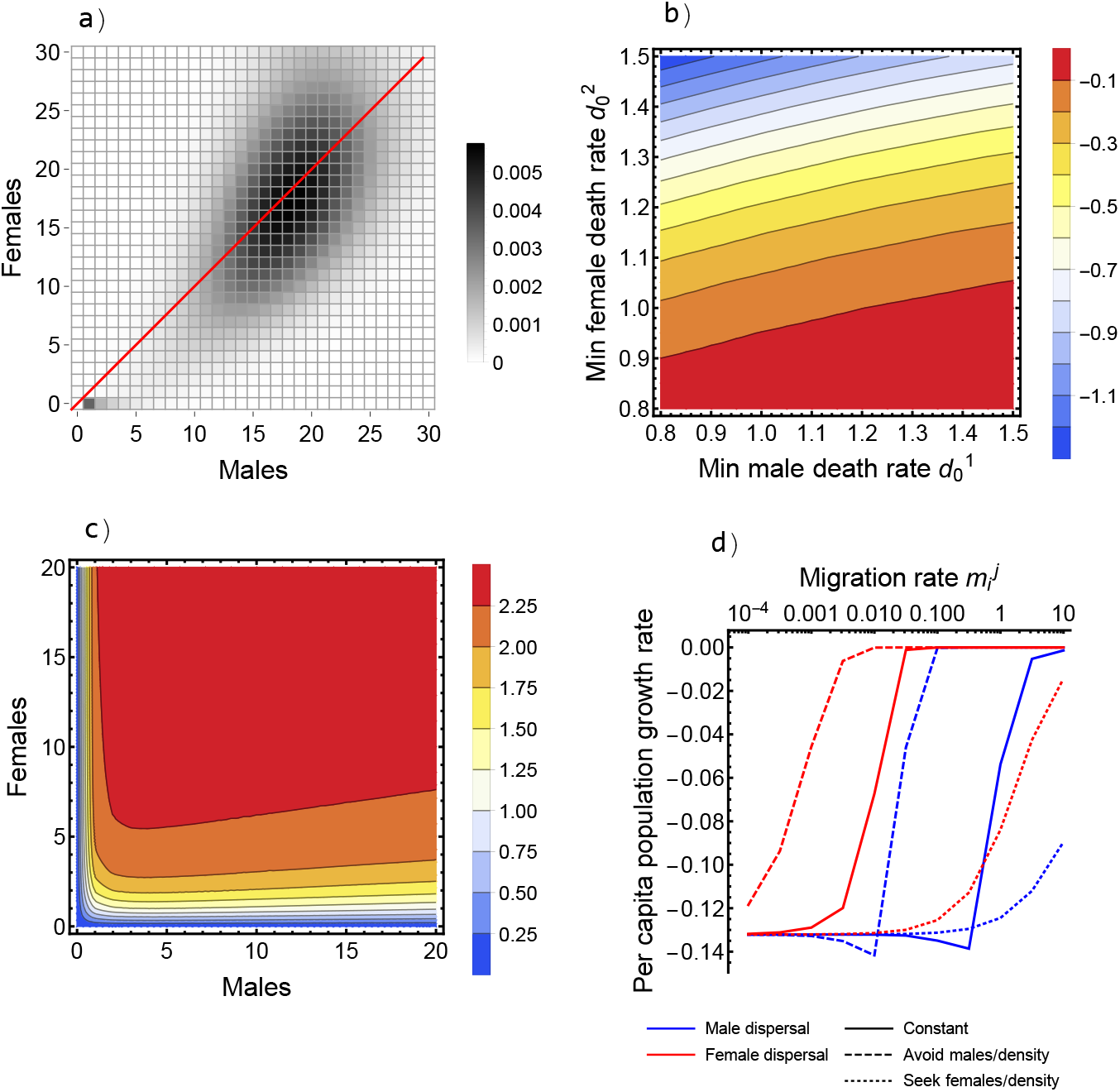
Results from the model with sex structure. (a) The group size distribution shows that group composition is less sensitive to the number of males in the group. (b) Per capita population growth rate (red is higher; see legend) increases with lower minimum female death rate 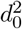 and higher minimum male death rate 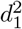 (up to a point). (c) Reproductive value (red is higher; see legend) is more sensitive to the number of females rather than the number of males in the group. (d) All forms of migration increase the per capita population growth rate, but female dispersal has a larger effect. Unless otherwise specified: *b*_0_ = 6, 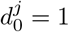, *d*_1_ = 0.05.

#### 5.3.2 Case study IV: Sex-biased migration

We now consider the consequences of sex-biased dispersal, motivated by the fact that males tend to be philopatric (and females disperse) in birds whereas females tend to be philopatric (and males disperse) in mammals (Greenwood, 1980). We define *m*_1_(*x*_1_, *x*_2_) to be the overall rate at which males migrate out of a group with *x*_1_ males and *x*_2_ females. We consider three dispersal strategies such that

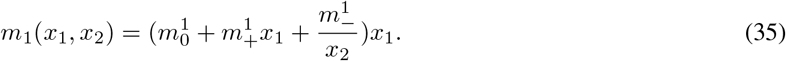

Here, 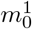 is a constant per capita male emigration rate, 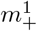 indicates a dispersal strategy where males seek to avoid groups with other males, and 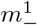 indicates a dispersal strategy where males seek to avoid groups with fewer females. Analogously, we consider female dispersal by

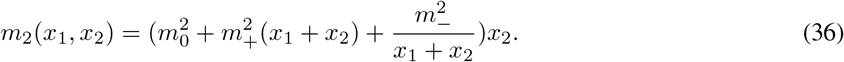

Now, 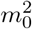 is a constant per capita female emigration rate, 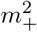 indicates a dispersal strategy where females seek to avoid groups with higher density, and 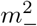 indicates a dispersal strategy where females seek to avoid groups with lower density. In practice, we consider only a single dispersal strategy at a time (setting all 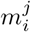 except one equal to 0) to isolate its impact on dynamics.

In general, when we add sex-biased dispersal to the model presented in the previous section (5.3.1), we find that all forms of dispersal increase the population’s per capita growth rate (Fig. 12d) for the reasons described in the section on migration in the model with ecologically identical individuals (Fig. 3). Female dispersal tends to provide a larger benefit than male dispersal (Fig. 12d). This occurs because group growth is largely decoupled from the number of males in the group once there is at least one male in the group. Dispersal strategies that involve seeking low density groups or avoiding males have a larger benefit to population growth than constant per capita migration, because this more efficiently redistributes individuals from large groups to small groups. In contrast, dispersal strategies that involve faster rates of emigration from small groups than large groups provide a smaller increase in the population growth rate (Fig. 12d).

### 5.4 Class-structured groups

Finally, we consider a case study where individuals belong to different classes in the social groups. We will use the example of eusocial insects with queens as type 1 individuals and workers as type 2 individuals. We note that a similar model form could be used to consider less extreme cases of division of labor such as dominant and subordinant individuals in cooperative breeders.

#### 5.4.1 Case study V: Division of labor

To keep the number of variables reasonable, we assume that each colony can have at most two queens. Reproduction can only occur if a colony has a queen, but birth rates increase (to saturation) as the colony’s number of workers increases. Specifically, the birth rate of queens is

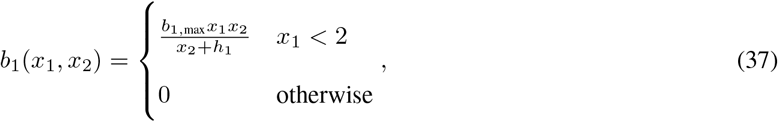

where *b*_1,max_ is the maximum queen birth rate and *h*_1_ is the number of workers that result in queens being born at rate *b*_1,max_/2. Worker birth rates follow the same form and are

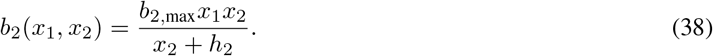

Queen death rates are assumed to decrease with the number of workers (due to e.g. better colony thermal regulation; Free and Spencer-Booth, 1958), such that

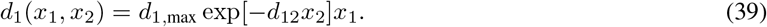

As with Eq. 31, this implies that the maximum death rate of each queen is *d*_1,max_ and *d*_12_ accounts for declining queen death rates with more workers. Worker death rates, on the other hand, increase with more workers due to competition within the colony. We assume worker death rates follow the same form of logistic death rates as Eq. 30,

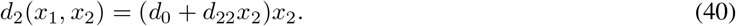

The key difference in the version of the model motivated by eusocial insects is the formation of new groups. We assume that colonies “swarm” and form new groups only if there are two queens in the group. Then, the formation of new groups occurs at rate *f*_+_*x*_2_. The number of workers that leaves with the new queen to form a new colony is given by the binomial PMF.

We find that the group size distribution consists of two types of groups. The first are groups where there is a single queen and the worker population is approximately at carrying capacity (orange line; Fig. 13a). The second are groups where there are no queens and the group is going extinct due to the lack of queen to produce more workers (blue line; Fig. 13a). If groups form more slowly (low *f*_+_), it is also possible to have groups with an excess of workers above the single-queen carrying capacity and two queens. We find that the per capita population growth rate is the lowest at intermediate queen birth rates and high worker birth rates (Fig. 13b). However, the average group size is largest when the worker birth rate is high and the queen birth rate is low (Fig. 13c). This is because colonies fission after the birth of a second queen and if queen birth occurs too frequently, then colonies do not have sufficient time to recover their worker population leading to small colonies with high extinction risk. Workers, on the other hand, ensure that the colony is able to continue to reproducing. The rate of change in the number of groups typically decreases with increasing queen birth rate (though a slight increase is seen at high queen birth rate due to many colonies forming via fissions; Fig. 13d). Increasing queen birth rate is generally detrimental for the colony (i.e., leads to the most rapid loss of groups). Nevertheless, intermediate queen birth rates still have the lowest rates of per capita population growth, because losing a single group has a larger influence on population growth rate when groups are larger, which occurs when queen birth rates are low. Though it may seem paradoxical, the fact that low queen birth rates are beneficial is related to the well-known separation in aging/lifespan timescales between queens and workers in eusocial insects (Keller and Genoud, 1997).

**Figure 13:**
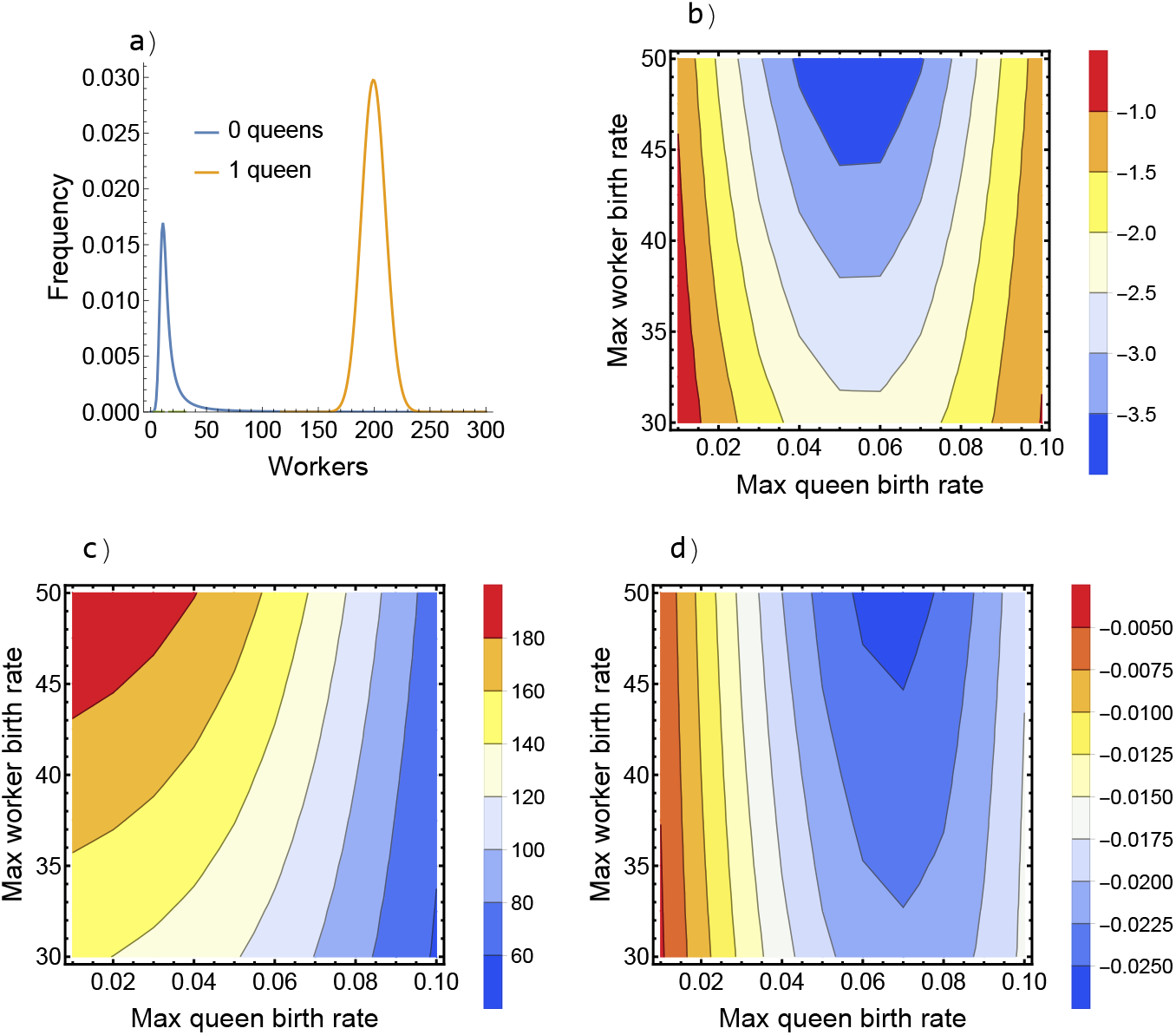
Results from the model with class structure. (a) The group size distribution shows groups with one queen and a carrying capacity number of workers (orange) and no queens that are going extinct (blue). (b) Per capita population growth rate is lowest at intermediate maximum queen birth rate *b*_1,max_. (c) The average group size is the largest with high worker birth rate and low queen birth rate. (d) The rate of change in number of groups primarily decreases with increasing queen birth rate. Unless otherwise specified: *b*_1,max_ = 0.05, *h*_1_ = 150, *b*_2,max_ = 30, *h*_2_ = 50, *d*_1,max_ = 0.003, *d*_12_ = 0.01, *d*_0_ = 0.01, *d*_22_ = 0.00055, *f*_+_ = 0.3.

## 6 Discussion

We have provided a general theoretical framework for studying the population dynamics of social species that is capable of capturing a wide variety of (potentially nonlinear) between-group processes, flexible forms of density dependence within social groups, and the fact that individuals can play distinct ecological roles. We developed multiple mathematical forms following this framework that each give different pieces of information about a system’s dynamics. We studied a series of between-group processes in isolation and considered interactions in special cases such as fission and fusions as well as fission and group competition. Through a series of case studies, we showed the importance of age, sex, and class structure in social populations while simultaneously demonstrating the applicability of our framework for capturing population dynamics in real systems. Below we 1) generalize the influence of each between-group process to provide a set of rules to determine how any process will influence dynamics, 2) discuss the role of density-dependence within social groups and describe how density dependence at the population level may emerge from between-group processes, 3) relate our results to past work on the importance of group size heterogeneity, 4) consider the analogy that groups are individuals to the population in the context of population ecology, 5) discuss the implications of our results for understanding social evolution, and 6) describe potential future directions for the study of the dynamics of socially structured populations.

### 6.1 A general procedure for understanding dynamics

By studying a series of between-group processes, we are able to delineate a set of general rules for how any between-group process can be expected to influence the population dynamics of a social species. The direct influence of a between-group process is to 1) change the number of groups and/or 2) alter the group size distribution. Between-group processes that increase the number of groups in the population (e.g., fissions) tend to increase the per capita population growth rate, whereas processes that decrease the number of groups (e.g., fusions, extinctions) tend to decrease the per capita population growth rate. The shift in the group size distribution can be thought of as a proximate cause of changes in population dynamics. Between-group processes that shift the group size distribution larger tend to increase the per capita population growth rate, since this decreases the rate that any one group can be expected to go extinct. Small groups (i.e., those well below carrying capacity) are uniquely important for understanding the population dynamics of social species since they are simultaneously at a heightened extinction risk due to demographic stochasticity and have the greatest potential for sustained growth back to their carrying capacity (provided they are above the Allee threshold, if one exists). This dual nature of small groups also helps to clarify the influence of various between-group processes. Broadly, when between-group processes have multiple effects, changing the number of groups plays a larger role in determining dynamics than shifting the group size distribution (as can be seen in the case of fissions and fusions).

### 6.2 The role and emergence of density dependence across scales

Our model recapitulates the result that within-group density dependence should not be expected to scale to the population level (Bateman et al., 2011; Angulo et al., 2013; Lerch et al., 2018; Bateman et al., 2018). Together with Bateman et al. (2018), we have shown that the null expectation (considering only within-group density dependence and potentially fissions/new group formation) should be that social populations will shrink or grow exponentially. Put simply, population-level density dependence requires that vital rates somehow change as a function of population size (or the number of groups) and not only as a function of group size. Due to this dependence on the group level, population-level density dependence in social species can arise in ways that are contingent upon between-group processes. Perhaps the most straightforward manifestation of this is the case in which group interaction rates increase with the number of groups (e.g., following a mass action assumption at the group level), leading to higher death rates in larger populations due to increased group competition (Fig. 9). Here, the group level mediates the effect of population size on individuals. More surprisingly, population-level density dependence can arise even if increasing population sizes do not directly lead to stronger competition. As an example, if fissions are coupled with fusions that become more frequent with more groups in the population, then logistic growth emerges at the population level (Fig. 8). In this case, the fact that the number of groups grows logistically directly translates into the number of individuals growing logistically: between-group processes alone are driving population-level density dependence. Notably, the form of density dependence that emerges at the population level is completely independent from the form of density dependence occurring within social groups.

All of this is not meant to imply that within-group density dependence is unimportant, simply that it does not determine the qualitative features of population-level density dependence. Within-group density dependence does have a large impact on the rate at which populations grow exponentially: within-group density dependence leading to heightened extinction risks for a single group (such as an Allee effect) results in more negative (or less positive) per capita population growth rates. Further, within-group density dependence has a large impact on the conditions that individuals are expected to experience (i.e., the average per capita group growth rate). Logistic growth and Allee effects often result in opposite trends for how a between-group process alters the average per capita group growth rate, potentially leading to dramatic consequences for individual fitness (discussed below).

### 6.3 The role of group size heterogeneity

Our results suggest that heterogeneity in group sizes is not a requirement to prevent within-group density dependence from scaling to the population level. This stands in contrast to a recent model (Angulo et al., 2018) that uses environmental and demographic stochasticity to generate heterogeneity in group size and also interactions at the group level. Patterns of stochasticity do drive emergent population-level effects (Angulo et al., 2018), but care must be taken in interpreting these results. Stochastic effects (*censu* Lande, 1998) likely play some role in addition to the intrinsic effects of the group size heterogeneity generated by stochasticity. Rather than relying on stochasticity to generate group size heterogeneity, our two variable model allows us to isolate the effects of heterogeneity *per se*. Consistent with Bateman et al. (2018), our results suggest that within-group dynamics alone never result in density-dependence at the population level. Group size heterogeneity may alter specific vital rates, but does not qualitatively alter outcomes, except in the small number of situations where changing the size of a specific group drove the population-level results (e.g. when fissions interact with intergroup competition). We note that in nature group size heterogeneity is likely to have important effects (we use it here to set the rate at which groups are lost from the model without group size heterogeneity), but that it is not a requirement for either a lack of scaling of within-group density dependence to the population level or to explain many patterns of population growth.

### 6.4 Groups are individuals to the population

A common metaphor in evolutionary biology is that groups themselves behave as organisms (or individuals) in social populations (Wilson and Sober, 1989; Clejan et al., 2022). Detractors argue that genetic variation and class structure make this a poor analogy (Gardner and Grafen, 2009; Gardner, 2015). Our results suggest that viewing groups as individuals to the population may be a valuable perspective for population ecologists to take, especially since they often ask questions that do not rely upon understanding the fitness consequences of sociality (and thus genetic relatedness). A number of results can be viewed through this lens. Fissions increase the per capita population growth rate because they correspond to the “birth” of new groups (in contrast to fusions that are the “death” of groups by decreasing the total number of groups by 1). Thus, it makes sense that if the fission rate increases linearly with the number of groups (analogous to a constant per capita birth rate) and the fusion rate increases quadratically with the number of groups (analogous to the standard assumption of death rate for logistic growth), then logistic growth emerges at the population level. The same effect famously emerges from Levins’ patch-occupancy model for metapopulations (Levins, 1969), in which quadratic patch colonization and linear patch extinction rates combine to make the proportion of occupied patches grow according to the logistic model exactly. Our work extends this result substantially, by showing it is robust to the inclusion of within-group dynamics and more complex mechanisms for creating and destroying groups.

Population extinction can also benefit from this perspective. Here, each group going extinct is analogous to deaths in the well-mixed case and thus extinction occurs only once the last group has gone extinct. Hence, increasing the extinction rate of a single group by, for example, decreasing its carrying capacity has an out-sized influence on population extinction risk (analogous to increasing individual death rates). Again, this perspective has connections to spatially-structured populations. For example, fragmentation raises extinction risk (independent of habitat loss) by lowering the subpopulation carrying capacity (Day and Possingham, 1995; Hanski, 1999; Crooks et al., 2017).

These results also relate to the dynamics of clonal plants (Eriksson, 1993). Similar models have been developed that treat clones (genets) as a single individual comprised of distinct parts (ramets) (Eriksson, 1994). We can think of genets as comparable to groups in our model whose dynamics play out on longer timescales and ramets as individual organisms.

### 6.5 Implications for evolution

As explained above, the model presented here was developed as a general theory of group selection (Simon, 2010; Simon et al., 2013). Given that so much work on social populations has been carried out on understanding the evolution of their cooperative behaviors (Nowak, 2006; Lehmann et al., 2006; Akçay, 2020) rather than their ecological dynamics, it is worth determining whether these results have implications for the study of social evolution. Our results provide a population ecologist’s perspective on a number of results from social evolution theory. First, we find that between-group processes often have opposing trends on the per capita population growth rate and the average per capita group growth rate. Since per capita group growth rate is a fitness proxy in social species, this would create tension between individual fitness and population growth, if behaviors mediating the frequency of between-group processes were under selection. This relates to previous work showing that the evolution of behavior and population dynamics are inherently unstable when considered jointly in social species (Halloway et al., 2020). Second, Allee effects, group size, and generally population demography is known to be important for sociality to evolve (Aviles, 1999; Avilés et al., 2002; Lehmann et al., 2006; Hauert et al., 2006; Lehmann and Rousset, 2010; van Veelen et al., 2010; Garcia and De Monte, 2013; Lerch and Abbott, 2020). However, the actual influence many of these features have on dynamics had not been clarified for social populations. Thus, the importance of population demography for social evolution could not be firmly grounded in population ecology. Third, behaviors underlying between-group processes should be expected to evolve. Since between-group processes shape the population’s social structure, their evolution will feedback with the evolution of cooperative behaviors. Such coevolutionary dynamics have been argued to be the next frontier for studying social evolution (Akçay, 2020). Relating such work to population ecology outlined here will allow for a clear assessment of the ecological side of these feedbacks.

### 6.6 Future directions

Though we have attempted to extend and reinterpret Simon et al.’s (2013) modeling framework to generate a comprehensive overview of the dynamics of socially-structured populations, there is of course always more investigation possible. We see two ways, in particular, that this framework could be useful for deepening our understanding of the dynamics of social populations: 1) using a wider variety of functional forms and 2) applying the model to specific study systems. First, we have mostly limited ourselves to studying the simplest functional forms for the rates of various between-processes (i.e., constant per capita or per group rates). Future theoretical work could characterize how sensitive our conclusions are to various functional forms. Inspiration for specific functional forms connects to our second, broad future direction, applying the model to real systems. In cases where manipulative experiments cannot be conducted, our modeling framework provides a flexible tool for understanding the dynamics of social populations. The model could be parameterized to match real populations to provide insight into specific species. Our case studies provide a blueprint for connecting our model to real systems. As a specific example, our case studies on juvenile group formation and class-structured groups demonstrate the importance of the mechanisms behind new group formation (which, for example, can lead to population-level demographic Allee effects; section 5.2.2). Mechanistically modeling and comparing a multitude of group formation strategies could be one way that this modeling framework could provide real-world insight into specific systems.

The most notable missing feature from our model involves properly accounting for individual behavior and collective decisions. Much work on social species focuses not broadly on their population dynamics, as we have, but rather on the dynamics and influence of their social networks (Krause et al., 2009; Farine and Whitehead, 2015). Integrating the population ecologists’ perspective on population dynamics with animal social networks provides a major goal for theoreticians interested in the study of social species. It is well-known that an individual’s position in their social network has major influence over their life trajectory. This is true in cooperative breeders with high reproductive skew, where dominant individuals have the highest fitness, such as elephant seals (*Mirounga* sp. Hoezel et al., 1999), pied babblers (*Turdoides bicolor*; Nelson-Flower et al., 2011), and meerkats (Griffin et al., 2003) and also in more egalitarian societies such as baboons (Silk et al., 2003, 2010; Archie et al., 2014), bottlenose dolphins (*Tursiops* sp.; Stanton and Mann, 2012), spotted hyenas (*Crocuta crocuta*; Strauss and Holekamp, 2019; Turner et al., 2021), and greater anis (*Crotophaga major*; Riehl and Strong, 2018). Further, social networks may alter the outcome of some between-group processes and individual behavioral decisions determine the structure of social networks following between-group processes (for example, in the case of fissions; Lerch et al., 2021). In addition to generating interesting feedbacks, properly accounting for individual behavior would begin to unify to largely disparate perspectives on social populations. Thus, in order to create a truly general, universally applicable framework for social species the roles of social networks, individual behavior, and collective decisions on populations dynamics must be clarified.

## 7 Acknowledgements

We thank members of the Servedio lab at UNC Chapel Hill and the Alberts lab at Duke University for feedback on the framing, model, and results, Sorin Mitran for discussions on numerical solutions of PDEs, and Liz Lange, Robin Snyder, and Matthew Zipple for helpful comments on an early draft of the manuscript. UNC’s research computing infrastructure supported our analysis.

## Notes

https://zenodo.org/record/7063010#.Yx8Q0aHMJ3i

### Competing Interest Statement

The authors have declared no competing interest.

## References

Erol Akçay. Deconstructing evolutionary game theory: Coevolution of social behaviors with their evolutionary setting. The American Naturalist, 195(2):315–330, 2020. doi: 10.1086/706811.

Elena Angulo, Greg S A Rasmussen, David W Macdonald, and Courchamp Franck. Do social groups prevent allee effect related extinctions?: The case of wild dogs. Journal of Animal Ecology, 10:11, 2013. doi: https://doi.org/10.1186/1742-9994-10-11.

Elena Angulo, Gloria M. Luque, Stephen D. Gregory, John W. Wenzel, Carmen Bessa-Gomes, Ludek Berec, and Franck Courchamp. Allee effects in social species. Journal of Animal Ecology, 87(1):47–58, 2018. doi: https://doi.org/10.1111/1365-2656.12759.

Elizabeth A. Archie, Jenny Tung, Michael Clark, Jeanne Altmann, and Susan C. Alberts. Social affiliation matters: both same-sex and opposite-sex relationships predict survival in wild female baboons. Proceedings of the Royal Society B: Biological Sciences, 281(1793):20141261, 2014. doi: 10.1098/rspb.2014.1261.

Leticia Aviles. Cooperation and non-linear dynamics: An ecological perspective on the evolution of sociality. Evolutionary Ecology Research, 1:459–477, 1999.

Leticia Avilés and Paul Tufiño. Colony size and individual fitness in the social spider anelosimus eximius. The American Naturalist, 152(3):403–418, 1998. doi: 10.1086/286178.

Leticia Avilés, Patrick Abbot, and Asher D. Cutter. Population ecology, nonlinear dynamics, and social evolution. i. associations among nonrelatives. The American Naturalist, 159(2):115–127, 2002. doi: 10.1086/324792.

A. W. Bateman, T. Coulson, and T. H. Clutton-Brock. What do simple models reveal about the population dynamics of a cooperatively breeding species? Oikos, 120(5):787–794, 2011. doi: https://doi.org/10.1111/j.1600-0706.2010.18952.x.

A. W. Bateman, A. Ozgul, J. F. Nielsen, T. Coulson, and T. H. Clutton-Brock. Social structure mediates environmental effects on group size in an obligate cooperative breeder, suricata suricatta. Ecology, 94(3):587–597, 2013. doi: https://doi.org/10.1890/11-2122.1.

Andrew W. Bateman, Arpat Ozgul, Tim Coulson, and Tim H. Clutton-Brock. Density dependence in group dynamics of a highly social mongoose, suricata suricatta. Journal of Animal Ecology, 81(3):628–639, 2012. doi: https://doi.org/10.1111/j.1365-2656.2011.01934.x.

Andrew W. Bateman, Arpat Ozgul, Martin Krkošek, and Tim H. Clutton-Brock. Matrix models of hierarchical demography: Linking group- and population-level dynamics in cooperative breeders. The American Naturalist, 192 (2):188–203, 2018. doi: 10.1086/698217.

Dominik M. Behr, John W. McNutt, Arpat Ozgul, and Gabriele Cozzi. When to stay and when to leave? proximate causes of dispersal in an endangered social carnivore. Journal of Animal Ecology, 89(10):2356–2366, 2020. doi: https://doi.org/10.1111/1365-2656.13300.

James H. Brown and Astrid Kodric-Brown. Turnover rates in insular biogeography: Effect of immigration on extinction. Ecology, 58(2):445–449, 1977. doi: https://doi.org/10.2307/1935620.

Damien Caillaud, Winnie Eckardt, Veronica Vecellio, Felix Ndagijimana, Jean-Pierre Mucyo, Jean-Paul Hirwa, and Tara Stoinski. Violent encounters between social units hinder the growth of a high-density mountain gorilla population. Science Advances, 6(45):eaba0724, 2020. doi: 10.1126/sciadv.aba0724.

Hal Cawell. Matrix Population Models. Oxford University Press, Oxford, UK, 2006.

D.L. Cheney and R.M. Seyfarth. The influence of intergroup competition on the survival and reproduction of female vervet monkeys. Behavioral Ecology and Sociobiology, 21:375–386, 1987. doi: https://doi.org/10.1007/BF00299932.

Iuval Clejan, Christopher D. Congleton, and Brian A. Lerch. The emergence of group fitness. Evolution, 76(8): 1689–1705, 2022. doi: https://doi.org/10.1111/evo.14549.

Franck Courchamp, Tim Clutton-Brock, and Bryan Grenfell. Multipack dynamics and the allee effect in the african wild dog, lycaon pictus. Animal Conservation, 3(4):277–285, 2000. doi: https://doi.org/10.1111/j.1469-1795.2000.tb00113.x.

Bernard J. Crespi and Douglas Yanega. The definition of eusociality. Behavioral Ecology, 6(1):109–115, 03 1995. doi: 10.1093/beheco/6.1.109.

Kevin R. Crooks, Christopher L. Burdett, David M. Theobald, Sarah R. B. King, Moreno Di Marco, Carlo Rondinini, and Luigi Boitani. Quantification of habitat fragmentation reveals extinction risk in terrestrial mammals. Proceedings of the National Academy of Sciences, 114(29):7635–7640, 2017. doi: 10.1073/pnas.1705769114.

Leon Danon, Ashley P. Ford, Thomas House, Chris P. Jewell, Matt J. Keeling, Gareth O. Roberts, Joshua V. Ross, and Matthew C. Vernon. Networks and the epidemiology of infectious disease. Interdisciplinary Perspectives on Infectious Diseases, 2011:284909, 2011. doi: 10.1155/2011/284909.

J.R. Day and H.P. Possingham. A stochastic metapopulation model with variability in patch size and position. Theoretical Population Biology, 48(3):333–360, 1995. doi: https://doi.org/10.1006/tpbi.1995.1034.

Brian Dennis. Allee effects in stochastic populations. Oikos, 96(3):389–401, 2002. doi: https://doi.org/10.1034/j.1600-0706.2002.960301.x.

Richard Durrett. Essentials of Stochastic Processes. Springer, Cham, Switzerland, 2016.

Anders Eriksson, Federico Elías-Wolff, Bernhard Mehlig, and Andrea Manica. The emergence of the rescue effect from explicit within- and between-patch dynamics in a metapopulation. Proceedings of the Royal Society B: Biological Sciences, 281(1780):20133127, 2014. doi: 10.1098/rspb.2013.3127.

Ove Eriksson. Dynamics of genets in clonal plants. Trends in Ecology and Evolution, 8(9):313–316, 1993. doi: 10.1016/0169-5347(93)90237-J.

Ove Eriksson. Stochastic population dynamics of clonal plants: Numerical experiments with ramet and genet models. Ecological Research, 9(3):257–268, 1994. doi: https://doi.org/10.1007/BF02348412.

Damien R. Farine and Hal Whitehead. Constructing, conducting and interpreting animal social network analysis. Journal of Animal Ecology, 84(5):1144–1163, 2015. doi: https://doi.org/10.1111/1365-2656.12418.

J. B. Free and Y. Spencer-Booth. Observations on the Temperature Regulation and Food Consumption of Honeybees (Apis Mellifera). Journal of Experimental Biology, 35(4):930–937, 12 1958. doi: 10.1242/jeb.35.4.930.

Thomas Garcia and Silvia De Monte. Group formation and the evolution of sociality. Evolution, 67(1):131–141, 2013. doi: https://doi.org/10.1111/j.1558-5646.2012.01739.x.

A. Gardner. The genetical theory of multilevel selection. Journal of Evolutionary Biology, 28(2):305–319, 2015. doi: https://doi.org/10.1111/jeb.12566.

A. Gardner and A. Grafen. Capturing the superorganism: a formal theory of group adaptation. Journal of Evolutionary Biology, 22(4):659–671, 2009. doi: https://doi.org/10.1111/j.1420-9101.2008.01681.x.

Joanna Gascoigne, Ludek Berec, Stephen Gregory, and Franck Courchamp. Dangerously few liaisons: a review of mate-finding allee effects. Population Ecology, 51:355–372, 2009. doi: https://doi.org/10.1007/s10144-009-0146-4.

Daniel T. Gillespie. Exact stochastic simulation of coupled chemical reactions. The Journal of Physical Chemistry, 81(25):2340–2361, 1977.

Paul J. Greenwood. Mating systems, philopatry and dispersal in birds and mammals. Animal Behaviour, 28(4): 1140–1162, 1980. doi: https://doi.org/10.1016/S0003-3472(80)80103-5.

Ashleigh S. Griffin, Josephine M. Pemberton, Peter N. M. Brotherton, Grant McIlrath, David Gaynor, Ruth Kansky, Justin O’Riain, and Timothy H. Clutton-Brock. A genetic analysis of breeding success in the cooperative meerkat (Suricata suricatta). Behavioral Ecology, 14(4):472–480, 07 2003. doi: 10.1093/beheco/arg040.

Abdel H. Halloway, Margaret A. Malone, and Joel S. Brown. Unstable population dynamics in obligate co-operators. Theoretical Population Biology, 136:1–11, 2020. doi: https://doi.org/10.1016/j.tpb.2020.09.002.

Ilkka Hanski. Metapopulation Ecology. Oxford University Press, Oxford, UK, 1999.

Tara R. Harris. Between-group contest competition for food in a highly folivorous population of black and white colobus monkeys (*Colobus guereza*). Behavioral Ecology and Sociobiology, 61:316–329, 2006. doi: https://doi.org/10.1007/s00265-006-0261-6.

Christoph Hauert, Miranda Holmes, and Michael Doebeli. Evolutionary games and population dynamics: maintenance of cooperation in public goods games. Proceedings of the Royal Society B: Biological Sciences, 273(1600):2565–2571, 2006. doi: 10.1098/rspb.2006.3600.

Gil J. B. Henriques, Burton Simon, Yaroslav Ispolatov, and Michael Doebeli. Acculturation drives the evolution of intergroup conflict. Proceedings of the National Academy of Sciences, 116(28):14089–14097, 2019. doi: 10.1073/pnas.1810404116.

A. Rus Hoezel, Burney J. Le Boeuf, Joanne Reiter, and Claudio Campagna. Alpha-male paternity in elephant seals. Behavioral Ecology and Sociobiology, 46:298–306, 1999. doi: 10.1007/s002650050623.

Rufus A. Johnstone, Michael A. Cant, Dominic Cram, and Faye J. Thompson. Exploitative leaders incite intergroup warfare in a social mammal. Proceedings of the National Academy of Sciences, 117(47):29759–29766, 2020. doi: 10.1073/pnas.2003745117.

Peter M. Kappeler. Sex roles and adult sex ratios: insights from mammalian biology and consequences for primate behaviour. Philosophical Transactions of the Royal Society B: Biological Sciences, 372(1729):20160321, 2017. doi: 10.1098/rstb.2016.0321.

L. Keller and M. Genoud. Extraordinary lifespans in ants: a test of evolutionary theories of ageing. Nature, 389: 958–960, 1997. doi: 10.1038/40130.

Oded Keynan and Amanda R. Ridley. Component, group and demographic allee effects in a cooperatively breeding bird species, the arabian babbler (*Turdoides squamiceps*). Oecologia, 182:153–161, 2016. doi: 10.1007/s00442-016-3656-8.

Timothy Killingback and Michael Doebeli. Spatial evolutionary game theory: Hawks and doves revisited. Proceedings of the Royal Society of London. Series B: Biological Sciences, 263(1374):1135–1144, 1996. doi: 10.1098/rspb.1996.0166.

J. Krause, D. Lusseau, and R James. Animal social networks: An introduction. Behavioral Ecology and Sociobiology, 63:967–973, 2009. doi: 10.1007/s00265-009-0747-0.

Thomas G. Kurtz. Solutions of ordinary differential equations as limits of pure jump markov processes. Journal of Applied Probability, 7(1):49–58, 1970. doi: 10.2307/3212147.

Russell Lande. Demographic stochasticity and allee effect on a scale with isotropic noise. Oikos, 83:353–358, 1998. doi: https://doi.org/3546849.

Laurent Lehmann and François Rousset. How life history and demography promote or inhibit the evolution of helping behaviours. Philosophical Transactions of the Royal Society B: Biological Sciences, 365(1553):2599–2617, 2010. doi: 10.1098/rstb.2010.0138.

Laurent Lehmann, Nicolas Perrin, and Francois Rousset. Population demography and the evolution of helping behaviors. Evolution, 60(6):1137–1151, 2006. doi: https://doi.org/10.1111/j.0014-3820.2006.tb01193.x.

Matthew A. Leibold and Jonathan M. Chase. Metacommunity Ecology. Princeton University Press, Princeton, NJ, USA, 2017.

Sylvain Lemoine, Anna Preis, Liran Samuni, Christophe Boesch, Catherine Crockford, and Roman M. Wittig. Between-group competition impacts reproductive success in wild chimpanzees. Current Biology, 30(2):312–318.e3, 2020. doi: https://doi.org/10.1016/j.cub.2019.11.039.

Brian A Lerch and Karen C Abbott. Allee effects drive the coevolution of cooperation and group size in high reproductive skew groups. Behavioral Ecology, 31(3):661–671, 2020. doi: 10.1093/beheco/araa009.

Brian A. Lerch, Ben C. Nolting, and Karen C. Abbott. Why are demographic allee effects so rarely seen in social animals? Journal of Animal Ecology, 87(6):1547–1559, 2018.

Brian A. Lerch, Karen C. Abbott, Elizabeth A. Archie, and Susan C. Alberts. Better baboon break-ups: collective decision theory of complex social network fissions. Proceedings of the Royal Society B: Biological Sciences, 288 (1964):20212060, 2021. doi: 10.1098/rspb.2021.2060.

Richard Levins. Some Demographic and Genetic Consequences of Environmental Heterogeneity for Biological Control. Bulletin of the Entomological Society of America, 15(3):237–240, 1969. doi: 10.1093/besa/15.3.237.

Dieter Lukas and Tim Clutton-Brock. Life histories and the evolution of cooperative breeding in mammals. Proceedings of the Royal Society B: Biological Sciences, 279(1744):4065–4070, 2012. doi: 10.1098/rspb.2012.1433.

Nino Maag, Gabriele Cozzi, Tim Clutton-Brock, and Arpat Ozgul. Density-dependent dispersal strategies in a cooperative breeder. Ecology, 99(9):1932–1941, 2018. doi: https://doi.org/10.1002/ecy.2433.

Bonaventura Majolo, Aurora deBortoli Vizioli, Laura Martínez-Íñigo, and Julia Lehmann. Effect of group size and individual characteristics on intergroup encounters in primates. International Journal of Primatology, 41:325–341, 2020. doi: https://doi.org/10.1007/s10764-019-00119-5.

A. Catherine Markham, Susan C. Alberts, and Jeanne Altmann. Intergroup conflict: ecological predictors of winning and consequences of defeat in a wild primate population. Animal Behaviour, 84(2):399–403, 2012. doi: https://doi.org/10.1016/j.anbehav.2012.05.009.

A. Catherine Markham, Laurence R. Gesquiere, Susan C. Alberts, and Jeanne Altmann. Optimal group size in a highly social mammal. Proceedings of the National Academy of Sciences, 112(48):14882–14887, 2015. doi: 10.1073/pnas.1517794112.

J. Maynard Smith. What use is sex? Journal of Theoretical Biology, 30(2):319–335, 1971. doi: https://doi.org/10.1016/0022-5193(71)90058-0.

J. Weldon McNutt. Sex-biased dispersal in african wild dogs,lycaon pictus. Animal Behaviour, 52(6):1067–1077, 1996. doi: https://doi.org/10.1006/anbe.1996.0254.

Jane Molofsky, Richard Durrett, Jonathan Dushoff, David Griffeath, and Simon Levin. Local frequency dependence and global coexistence. Theoretical Population Biology, 55(3):270–282, 1999. doi: https://doi.org/10.1006/tpbi.1998.1404.

Martha J. Nelson-Flower, Phil A.R. Hockey, Colleen O’Ryan, Nichola J. Raihani, Morné A. du Plessis, and Amanda R. Ridley. Monogamous dominant pairs monopolize reproduction in the cooperatively breeding pied babbler. Behavioral Ecology, 22(3):559–565, 03 2011. doi: 10.1093/beheco/arr018.

M. Nowak and R. May. Evolutionary games and spatial chaos. Nature, 359(6398):826–829, 1992. doi: 10.1038/359826a0.

Martin A. Nowak. Five rules for the evolution of cooperation. Science, 314(5805):1560–1563, 2006. doi: 10.1126/science.1133755.

Craig Packer, Ray Hilborn, Anna Mosser, Bernard Kissui, Markus Borner, Grant Hopcraft, John Wilmshurst, Simon Mduma, and Anthony R. E. Sinclair. Ecological change, group territoriality, and population dynamics in serengeti lions. Science, 307(5708):390–393, 2005. doi: 10.1126/science.1105122.

Danai Papageorgiou and Damien R. Farine. Group size and composition influence collective movement in a highly social terrestrial bird. eLife, page e59902, 2020. doi: 10.7554/eLife.59902.

Anatolii Puhalskii and Burton Simon. Discrete evolutionary birth-death processes and their large population limits. Stochastic Models, 28(3):388–412, 2012. doi: 10.1080/15326349.2012.699752.

Gregory S. A. Rasmussen, Markus Gusset, Franck Courchamp, and David W. Macdonald. Achilles’ heel of sociality revealed by energetic poverty trap in cursorial hunters. The American Naturalist, 172(4):508–518, 2008. doi: 10.1086/590965.

Christina Riehl and Meghan J. Strong. Stable social relationships between unrelated females increase individual fitness in a cooperative bird. Proceedings of the Royal Society B: Biological Sciences, 285(1876):20180130, 2018. doi: 10.1098/rspb.2018.0130.

Joan B. Silk, Susan C. Alberts, and Jeanne Altmann. Social bonds of female baboons enhance infant survival. Science, 302(5648):1231–1234, 2003. doi: 10.1126/science.1088580.

Joan B. Silk, Jacinta C. Beehner, Thore J. Bergman, Catherine Crockford, Anne L. Engh, Liza R. Moscovice, Roman M. Wittig, Robert M. Seyfarth, and Dorothy L. Cheney. Strong and consistent social bonds enhance the longevity of female baboons. Current Biology, 20(15):1359–1361, 2010. doi: https://doi.org/10.1016/j.cub.2010.05.067.

T. Scott Sillett, Nicholas L. Rodenhouse, and Richard T. Holmes. Experimentally reducing neighbor density affects reproduction and behavior of a migratory songbird. Ecology, 85(9):2467–2477, 2004. doi: https://doi.org/10.1890/03-0272.

Burton Simon. A dynamical model of two-level selection. Evolutionary Ecology Research, 12:555–588, 2010.

Burton Simon, Jeffrey A. Fletcher, and Michael Doebeli. Towards a general theory of group selection. Evolution, 67 (6):1561–1572, 2013.

M. J. Somers, J. A. Graf, M. Szykman, R. Slotow, and M. Gusset. Dynamics of a small re-introduced population of wild dogs over 25 years: Allee effects and the implications of sociality for endangered species’ recovery. Oecologia, 158(4):239–247, 2008. doi: 10.1007/s00442-008-1134-7.

M. A. Stanton and J. Mann. Early social networks predict survival in wild bottlenose dolphins. PLoS ONE, 7(10): e47508, 2012. doi: 10.1371/journal.pone.0047508.

Eli D. Strauss and Kay E. Holekamp. Social alliances improve rank and fitness in convention-based societies. Proceedings of the National Academy of Sciences, 116(18):8919–8924, 2019. doi: 10.1073/pnas.1810384116.

Patrick L. Thompson, Laura Melissa Guzman, Luc De Meester, Zsófia Horváth, Robert Ptacnik, Bram Vanschoen-winkel, Duarte S. Viana, and Jonathan M. Chase. A process-based metacommunity framework linking local and regional scale community ecology. Ecology Letters, 23(9):1314–1329, 2020. doi: https://doi.org/10.1111/ele.13568.

Audrey Trochet, Elodie A. Courtois, Virginie M. Stevens, Michel Baguette, Alexis Chaine, Dirk S. Schmeller, Jean Clobert, and John J. Wiens. Evolution of sex-biased dispersal. The Quarterly Review of Biology, 91(3):297–320, 2016. doi: 10.1086/688097.

Julie W. Turner, Alec L. Robitaille, Patrick S. Bills, and Kay E. Holekamp. Early-life relationships matter: Social position during early life predicts fitness among female spotted hyenas. Journal of Animal Ecology, 90(1):183–196, 2021. doi: https://doi.org/10.1111/1365-2656.13282.

Mitthijs van Veelen, Julian Garcia, and Leticia Aviles. It takes grouping and cooperation to get sociality. Journal of Theoretical Biology, 264(4):1240–1253, 2010. doi: 10.1016/j.jtbi.2010.02.043.

Simon van Vliet and Michael Doebeli. The role of multilevel selection in host microbiome evolution. Proceedings of the National Academy of Sciences, 116(41):20591–20597, 2019. doi: 10.1073/pnas.1909790116.

David Sloan Wilson and Elliott Sober. Reviving the superorganism. Journal of Theoretical Biology, 136(3):337–356, 1989. doi: https://doi.org/10.1016/S0022-5193(89)80169-9.

Rosie Woodroffe. Demography of a recovering African wild dog (Lycaon pictus) population. Journal of Mammalogy, 92(2):305–315, 2011.

Rosie Woodroffe, Helen M. K. O’Neill, and Daniella Rabaiotti. Within- and between-group dynamics in an obligate cooperative breeder. Journal of Animal Ecology, 89(2):530–540, 2020a. doi: https://doi.org/10.1111/1365-2656.13102.

Rosie Woodroffe, Daniella Rabaiotti, Dedan K. Ngatia, Thomas R. C. Smallwood, Stefanie Strebel, and Helen M. K. O’Neill. Dispersal behaviour of african wild dogs in kenya. African Journal of Ecology, 58(1):46–57, 2020b. doi: https://doi.org/10.1111/aje.12689.

